# Multiplexed and multivariate representations of sound identity during perceptual constancy

**DOI:** 10.1101/102889

**Authors:** Stephen M. Town, Katherine C. Wood, Jennifer K. Bizley

**Author notes:** Corresponding authors: Stephen Town Jennifer Bizley.

## Abstract

Perceptual constancy requires neural representations that are selective for object identity, but also tolerant for identity-preserving transformations. How such representations arise in the brain and contribute to perception remains unclear. Here we studied tolerant representations of sound identity in the auditory system by recording multi-unit activity in tonotopic auditory cortex of ferrets discriminating the identity of vowels which co-varied across orthogonal stimulus dimensions (fundamental frequency, sound level, location and voicing). We found that neural decoding of vowel identity was most successful across the same orthogonal dimensions over which animals generalized their behavior. We also decoded orthogonal sound features and behavioral variables including choice and accuracy to show a behaviorally-relevant, multivariate and multiplexed representation of sound, with each variable represented over a distinct time-course. Finally, information content and timing of sound feature encoding was modulated by task-engagement and training, suggesting that tolerant representations during perceptual constancy are attentionally and experience-dependent.

## Introduction

Perceptual constancy, also known as perceptual invariance, is the ability to recognize objects across variations in sensory input such as a face from multiple angles or a word spoken by different talkers (Bizley and Cohen, 2013; Logothetis and Sheinberg, 1996). Perceptual constancy requires that sensory systems such as vision and hearing develop a level of tolerance to identity preserving transformations (DiCarlo and Cox, 2007; DiCarlo et al., 2012). In hearing, the development of tolerance is critical to the representation of sounds such as individual words or phonemes across talkers, voice pitch, background noise and other acoustic transformations (Sharpee et al., 2011) and is a key step in auditory object formation and scene analysis (Bizley and Cohen, 2013; Bregman, 1990; Griffiths et al., 2004).

Both humans and other animals perceive sound features constantly despite variation in sensory input: we can recognize loudness across variation in location (Zahorik and Wightman, 2001), frequency across sound level (Polley et al., 2006) and sound identity across talkers (Kojima and Kiritani, 1989; Ohms et al., 2010), vocal tract length (Ghazanfar et al., 2007; Schebesch et al., 2010; Smith et al., 2005) and fundamental frequency (F0)(Bizley et al., 2013a; Honorof and Whalen, 2010; Town et al., 2015). At the neural level, tolerance emerges within auditory cortex for sounds including vocalizations (Billimoria et al., 2008; Carruthers et al., 2015; Meliza and Margoliash, 2012), pure tones (Sadagopan and Wang, 2008) and pulse trains (Bendor and Wang, 2007). Auditory cortical neurons are modulated by multiple features of speech sounds, such as synthesized vowels (Bizley et al., 2009), and when variables are considered in discrete time windows, tolerant responses of vowel identity, as well as information about sound location and F0 can be recovered (Walker et al., 2011). However, tolerance has yet to be shown in subjects actively demonstrating perceptual constancy, and the behavioral relevance of tolerant representations in auditory cortex remains unclear. Furthermore, although auditory cortical processing is modulated by attention and experience (Osmanski and Wang, 2015), it is unknown how these processes affect tolerant representations.

Here we asked if tolerant representations exist in auditory cortex during perceptual constancy, how tolerance was related to behavior, and modulated by attention and experience. To address these questions, we recorded the activity of auditory cortical neurons in ferrets discriminating synthesized vowel sounds across identity-preserving, orthogonal acoustic transformations - including variations in F0, sound location, level and voicing.

We hypothesised that auditory cortical neurons would show tolerance across the same range of orthogonal variables over which animals demonstrate perceptual constancy, and that such tolerance would be degraded in cases where animals failed to generalize vowel identity. As auditory cortex represents multiple stimulus variables, we expected tolerance to be accompanied by information about both task-relevant and irrelevant sound features. Finally, we predicted that the neural correlates of perceptual constancy should be dependent on an animal’s behavioral performance, attentional state and training. Our findings confirmed that neurons could represent vowel identity across orthogonal variations and thus provide tolerant representations in perceptual constancy. Furthermore, we also demonstrated these representations were sensitive to behavioral performance, failures to perceive vowel constancy, attentional state and long-term experience.

## Results

### Perceptual constancy during vowel discrimination

To establish a behavioral model of perceptual constancy, ferrets were trained in a two-choice task (Fig 1A) to identify synthesized vowels varying in F0 (149 – 459 Hz), location (±90°), sound level (45 – 82.5 dB SPL), or voicing (in which vowels were generated to sound whispered and presented on 20% of trials as probe trials). Changes in these task-irrelevant orthogonal dimensions produced different spectra while preserving the formants peaks in the spectral envelope (Fig 1B) critical for vowel identification (Peterson and Barney, 1952; Town and Bizley, 2013). On each trial, the animal visited a central port to trigger presentation of the stimulus: two tokens of *the same* vowel, each lasting 250 ms with an inter-stimulus interval of 250 ms. Subjects then responded at a left / right spout depending on vowel identity, with correct responses rewarded with water and errors leading to a brief timeout (1-5 s). In each test session, vowels varied across only one orthogonal dimension (i.e. F0, level, location or voicing). Variation in each orthogonal dimension was sufficient that had the animals been discriminating these acoustic features performance would have been at, or close to, ceiling (Hine et al., 1994; Sinnott et al., 1992; Walker et al., 2011; Walker et al., 2009; Wood et al., 2017).

Ferrets discriminated vowels accurately across orthogonal dimensions: Performance was consistent across F0s (Fig 1C) and across locations (Fig 1D) (no effect of orthogonal dimension, logistic regression, *p* > 0.05, Table S1) and significantly better than chance at each F0 and location tested (binomial test vs. 50%, *p* < 0.001, Table S2). For all sound levels, performance was also better than chance (Fig 1E and Table S2), however performance increased significantly (but moderately) with sound level in 3 / 4 ferrets (*p* < 0.01; Table S1). Nevertheless performance was constant over a range of intensities and performance at lowest sound levels still exceeded chance. In contrast to the other orthogonal dimensions, ferrets failed to generalize across voicing: performance was significantly worse for whispered than voiced vowels (Fig 1F) and only two ferrets discriminated whispered stimuli better than chance (Table S2). These results confirmed that ferrets perceived a constant vowel identity across variations in acoustic input related to F0, sound location and sound level but not voicing, and so we predicted that we would find tolerant representations of vowel identity across changes in F0, location and sound level.

**Figure 1.**
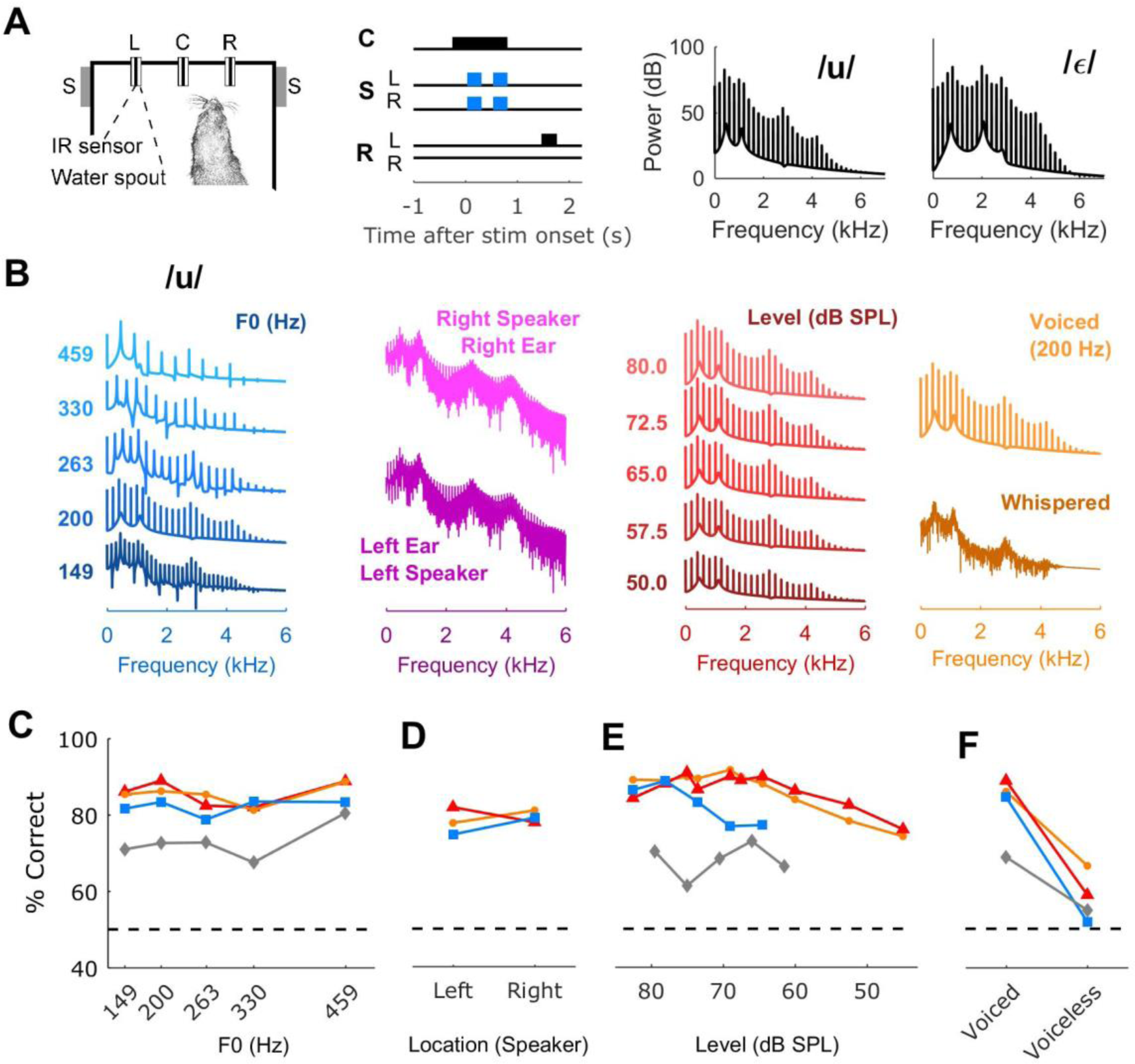
Perceptual constancy during vowel discrimination. **(A)** Schematic of task design: Animals initiated trials by visiting a central port (C) and waiting for a variable period before stimulus presentation. Speakers (S) presented sounds (two tokens *of the same* vowel; blue) to the left and right of the head in all conditions except when sound varied across location - in which case they were presented from either left (S_L_) or right (S_R_) speaker only. Animals responded at the left or right spout depending on vowel identity. **(B)** Spectra for 13 examples of one vowel /u/ with varying F0, location, sound level and voicing. Spectra for sounds across location were generated in virtual acoustic space (Schnupp et al., 2003) although sounds varied in free-field location. **(C-F)** Behavioral performance when discriminating vowels across F0 (C), location (D), level (E) and location (F). Individual subjects are shown as separate lines.

### Decoding sound features

We implanted arrays of independently moveable tungsten microelectrodes in left and right auditory cortex, where electrodes targeted the low frequency reversal between tonotopic primary and posterior fields (Bizley et al., 2005; See Fig S2 in Town et al., 2017). We recorded 471 sound-responsive multi-units and, for each unit, measured responses to vowels across F0, sound location, level and voicing during task performance (Fig 2A). We quantified the information available about vowel identity, F0, etc., by decoding stimulus features in one dimension across changes in the orthogonal dimension from single trial responses. Our decoder compared the Euclidean distances of time-varying patterns of neural activity, with leave-one-out cross validation (Foffani and Moxon, 2004)(Fig S1A). The time window over which responses were decoded was variable and we searched for those parameters (start time and window duration) that gave best decoding performance (Fig S1B). Optimization significantly improved decoding performance (Fig S2, rank-sum, *p* < 0.001) and enabled comparison of the time windows over which units were maximally informative. We decoded responses from correct trials only as we reasoned these would provide the clearest demonstration of auditory cortical encoding. For each unit, we reported decoding performance (Fig 2B) and whether the unit could be classified as significantly informative as determined by a permutation test (*p* < 0.05, Fig 2C and Fig S1C), indicating that the unit provided a tolerant representation of vowel identity across variation in sensory input resulting from changes in orthogonal dimensions.

We found that the proportion of vowel informative units was highest across dimensions over which behavioral performance was most constant. Across variation in F0, 42.1% of units (154/366) were informative about vowel identity, 43.5% (50/115) were informative across varying sound location and 40.6% (80/197) across varying sound level, whereas only 30.4% (63/207) were informative about vowel identity across voicing (Fig 2C). Furthermore, when we decoded vowel identity at each orthogonal value (Fig 2D) we found that the proportion of vowel informative units was independent of variation in F0 (logistic regression, χ^2^ = 0.776, *p* = 0.378), location (χ^2^ = 2.17, *p* = 0.140), level (χ^2^ = 0.447, *p* = 0.504) and voicing (χ^2^ = 0.983, *p* = 0.321). Together this suggests that auditory cortical neurons provide representations of vowel identity that are tolerant to variations in acoustic input caused by changes in F0, sound location and level, and to a lesser extent, voicing.

### Conserved information content

If units that represent vowel identity across one orthogonal dimension provide a truly tolerant representation, they should also represent vowel identity across multiple orthogonal dimensions. To test this, we counted the number of sound-responsive units from the entire recorded population that were vowel informative across F0, sound location, level and / or voicing.

While not every unit was tested across every orthogonal dimension, we found that units remained informative about vowel identity across the dimensions (F0, sound location and level) over which behavioral performance was constant (Fig 2E). Across sound level and location, 38.6% of units (22/57) were vowel informative across both dimensions. This value was close to the proportion of units sensitive to vowel identity across level or location (which were 40% and 43% respectively), indicating that the majority of vowel informative units represented sound identity across both orthogonal dimensions. Similarly, 34.8% of units (54/155) were informative about vowel identity across F0 and level, 34.4% of units (32/93) across F0 and location and 37.0% of units (20/54) were informative across all of F0, sound level and location. In contrast, notably fewer units (<22.5%) were informative about vowel identity over the other combinations of orthogonal factors – of which all included voicing, and across which, animals also generalized poorly. These findings suggest that a sizeable subpopulation of units provide tolerant information about vowel identity across orthogonal dimensions during perceptual constancy.

### Encoding of orthogonal dimensions

In addition to encoding vowel information during perceptual constancy, we also asked if neural responses conveyed information about orthogonal features of sounds that were irrelevant for task performance (Fig 2D). When considering all F0s or sound levels, we found 21.3% of units (78/366) were informative about F0 across vowels, and 24.9% (49/197) about sound level. These proportions increased to 38.5% (142/369) across F0; and 40.3% (29/72) across sound level when we decoded across the most extreme orthogonal values tested (149 vs 459 Hz or 45 vs 75 dB SPL). While a similar percentage of units (38.3%, 44/115) were informative about sound location, a greater proportion of units (50.7%, 105/207) were informative about voicing. Thus, the balance of units encoding task relevant and orthogonal dimensions was important for perceptual constancy: the proportion of units conveying information about vowel identity was greater than (F0, level) or similar to (location) the dimensions over which animals generalized, whereas across voicing, the balance of informative units was shifted towards the orthogonal dimension (50% to 30%; Fig. 2C).

We also tested whether units that were informative about one orthogonal variable (e.g. sound location) were also informative about other orthogonal variables (e.g. voicing). While we observed that some units were informative about multiple orthogonal dimensions (Fig 2F), such groups were significantly smaller than the corresponding analysis of vowel identity (sign-rank test on proportion of conserved units, *p* = 0.0098). Thus, while information about vowel identity was conserved *across* orthogonal dimensions, few units were informative about *multiple* orthogonal dimensions.

**Figure 2.**
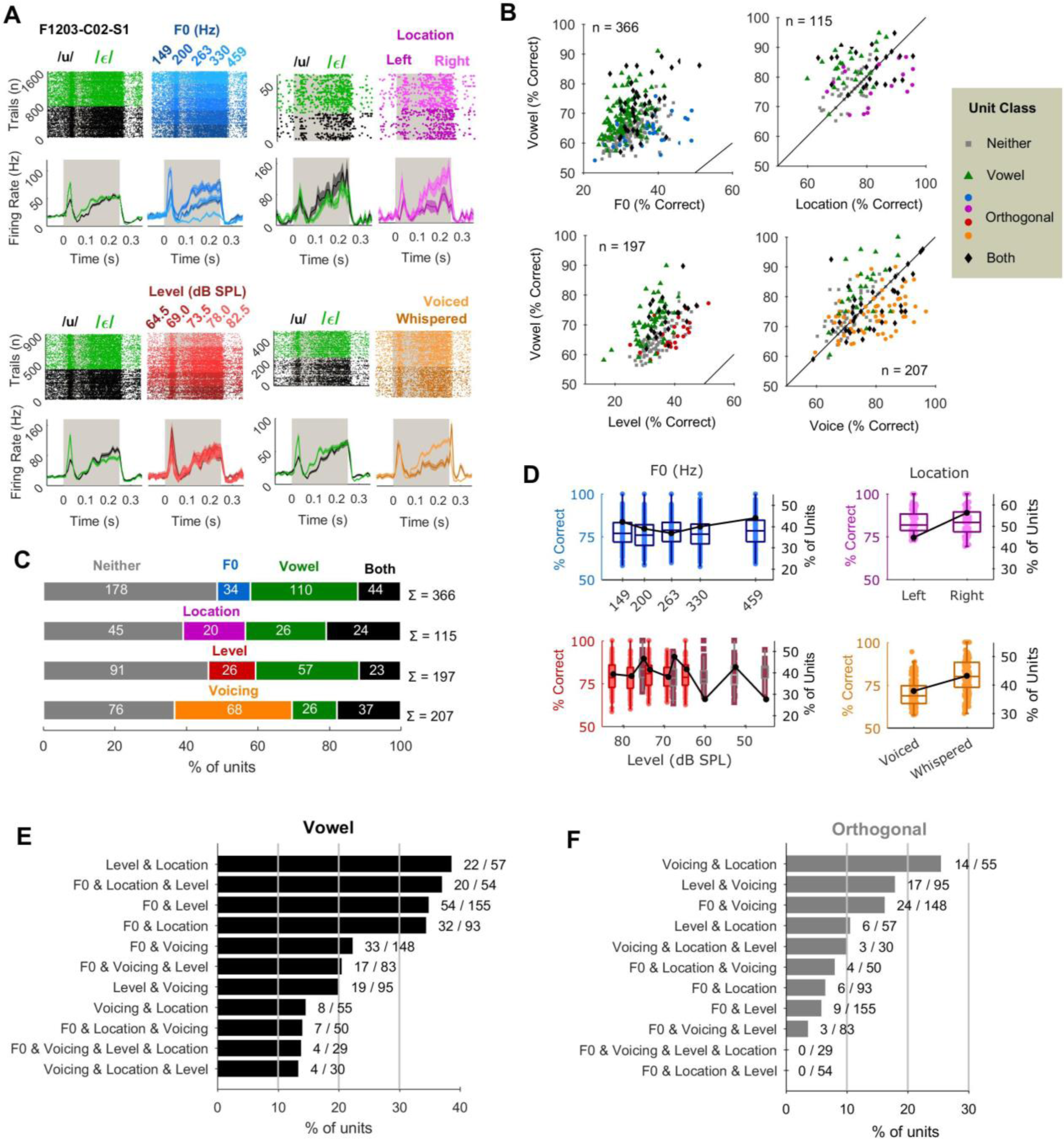
Neural responses and decoding acoustic features. **(A)** Raster and peri-stimulus time histograms (PSTHs) of neural responses of one unit to vowels across orthogonal variation in F0, location, level and voicing. Data plotted during presentation of the first sound token (grey bar) by vowel identity and by the orthogonal variable. PSTHs show mean ± s.e.m. firing rate. **(B)** Decoding performance when reconstructing vowel identity and orthogonal values from single trial responses of individual units. Data points indicate the best decoding performance of each unit. Chance performance for vowel identity, location and voicing was 50% and 20% for F0 and sound. (**C**) Number of units informative about vowel identity and / or orthogonal values when considering responses across all orthogonal values tested. **(D)** Decoder performance and proportion of vowel informative units at each orthogonal value. For sound level, different shades reflect the distinct sound ranges over which units were tested. **(E)** Number of units classified as vowel informative *across* multiple orthogonal dimensions. **(F)** As E but for units classified as being informative *about* multiple orthogonal dimensions.

### Temporal multiplexing of sound features

Our data show that multiple sound features are represented in auditory cortex, in some cases by the same neurons. Multivariate encoding in auditory cortex has been linked to temporal multiplexing, where units encode information about different stimulus features at distinct time points (Walker et al., 2011). We therefore asked whether multiplexing occurred during perceptual constancy and if the representation of vowel identity across orthogonal features was matched by conserved timing of information.

To study multiplexing, we compared the time windows that gave best performance decoding each stimulus feature following optimization. A time window was defined by its start time and duration, which we summarized as its midpoint (start time + duration/2). For each unit, we measured the midpoint for best decoding vowel identity across each orthogonal dimension, and for decoding each orthogonal dimension across vowel identity. We then compared cumulative distribution functions (CDFs) of midpoints across units that were informative about multiple stimulus features (dual-feature units) or only one feature (single-feature units).

We first confirmed the occurrence of multiplexing during perceptual constancy, finding that for dual-feature units, information about each feature emerged over a distinct time-course. Information about vowel identity arose significantly earlier than F0 (Fig 3A, Sign-rank test: z = -2.43, *p* = 0.015) and sound level (Fig 3C, z = -2.13, *p* = 0.033) whereas information about sound location arose significantly earlier than vowel identity (Fig 3B, z = 2.26, *p* = 0.024). In contrast, there was no significant difference in the timing of vowel identity and voicing (z = 1.33, *p* = 0.184). Thus multiplexing in these units only occurred for sounds that animals showed perceptual constancy.

For single-feature units, vowel identity was also best decoded earlier than F0 (Wilcoxon rank-sum test: z = -2.31, *p* = 0.021) but the differences between decoding of vowel identity and sound level (z = -0.933, *p* = 0.351), and vowel identity and location (z = 1.29, *p* = 0.198) were not significant. For single feature units, vowel identity was decoded significantly later than voicing (z =2.79, *p* = 0.005).

Across dual-feature units the timing of vowel information was conserved: CDFs did not differ significantly across orthogonal dimensions (Fig 3E, Kruskal-Wallis test: χ^2^ = 5.76, *p* = 0.124). In contrast, single-feature units showed significant differences in timing of vowel identity information across orthogonal dimensions (χ^2^ = 19.95, *p* = 1.74 x 10^-4^) with post-hoc comparisons showing that information about vowel identity across voicing emerged significantly later than across every other orthogonal factor (Tukey-Kramer corrected, F0: *p* = 0.001, location: *p* = 0.002, level: *p* = 0.013). Dual-feature units also encoded information about different orthogonal dimensions at significantly different times (Fig 3F, χ^2^ = 9.77, *p* = 0.012) with post-hoc comparisons showing F0 was decoded significantly later than location (*p* = 0.020) and voicing (*p* = 0.047). Similar results were also found for single feature units where encoding of orthogonal dimensions differed significantly in time (χ^2^ = 11.04, *p* = 0.0206) with post-hoc comparisons showing location was decoded significantly earlier than F0 (*p* = 0.026) and sound level ( *p* = 0.028).

These results emphasise an important role for temporal multiplexing: perceptual constancy only occurred when neurons that were sensitive to multiple stimulus features encoded information about each dimension in distinct time windows. Moreover, while the relative timing of vowel information and orthogonal dimensions was not important for generalisation, when vowel information was shifted in time, as in the case of voicing, perceptual constancy failed. Very similar results were observed when considering the start time or decoding window duration (Fig S4 and S5).

**Figure 3.**
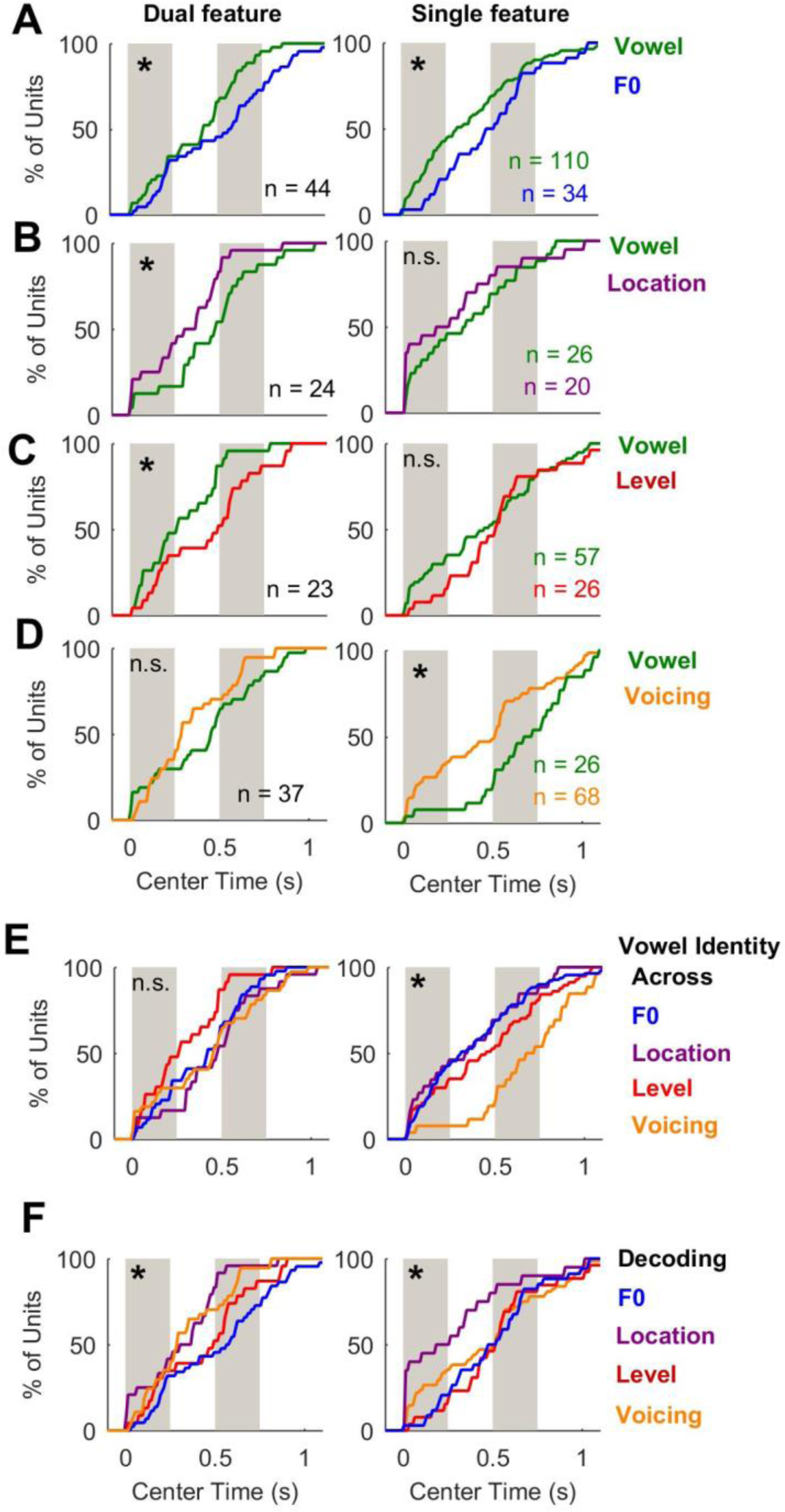
Temporal multiplexing in auditory cortex. **(A-D)** Cumulative distributions showing center times for best performance when decoding vowel identity or orthogonal variables (**A:** F0, **B**: location, **C**: level and **D**: voicing). Units are shown separately by classification as informative about vowel identity and orthogonal values (Dual feature units), or only vowel identity or orthogonal values (Single feature units). Grey bars represent the duration of each vowel token. **(E)** CDFs for decoding vowel identity across each orthogonal variable. **(F)** CDFs for decoding orthogonal values across vowels. Asterisks show significant differences between vowel and orthogonal (A-D, rank-sum or sign-rank depending on pairing, *p* < 0.05) or across orthogonal variables (Kruskal-Wallis, *p* < 0.05).

### Population decoding matches behavioral performance

Our results show a tolerant representation of vowel identity in auditory cortex during perceptual constancy; however we also wanted to understand how neural encoding was related to behavior. To match the subject’s behavioral performance, decoding performance must reach 100% as we only decoded neural responses on correct trials. While few individual units reached this level, it was possible to decode sound features with 100% performance from small populations of units (Fig 4A). Population decoding summed the number of individual units estimating each value of a stimulus feature (e.g. vowel /u/ or /ε/) with a weighting based on the relative spike-distance between decoding templates and test trials (see Methods). Decoding improved with population size, following a logistic function (Fig 4B, *p*<0.001) that allowed us to find the minimum number of units required reach 100% performance and compare decoding across conditions (logistic regression, analysis of deviance on main effect of stimulus feature). To equate the number of stimulus features decoded (n=2), we compared decoding across vowel, location and voicing (voiced and whispered) with the F0s (149 and 459 Hz) and sound levels (45 and 75 dB SPL) of greatest separation.

Population decoding required fewer units to perfectly reconstruct vowel identity across the orthogonal dimensions across which animals also showed perceptual constancy: To reach 100% performance across F0 required 15 units, and across sound location required 14 units whereas across sound level required 23 units and across voicing required 24 units (Fig 4B; see also Fig S3). Comparison of population growth curves confirmed that the growth of performance with population size was significantly different across the orthogonal dimensions across which vowel identity was decoded (χ^2^ = 315.3, *p* < 0.001). Vowel decoding performance increased significantly faster with population size than did decoding of F0 (χ^2^ = 319.7, *p* < 0.001), or equivalent to the orthogonal dimension (location, χ^2^ = 1.87, *p* = 0.172). Conversely, growth curves for population decoding of vowel identity rose significantly more slowly than for decoding of orthogonal features that animals’ failed to generalize across (voicing: χ^2^ = 174.9, *p* < 0.001) or incompletely generalised (sound level: χ^2^ = 40.9, *p* = 1.62 x 10^-10^). Population decoding of orthogonal values also differed significantly across dimensions (χ^2^ = 191.5, *p* < 0.001), with 23 units required to decode F0 with 100% performance, 13 units for sound location, 20 units for sound level with 20 units and 15 units for voicing. These findings, suggest that the dynamics of population decoding reflect the ability of animals to generalise: a hallmark of perceptual constancy across a given dimension is that a performance can be supported by a smaller number of units.

**Figure 4.**
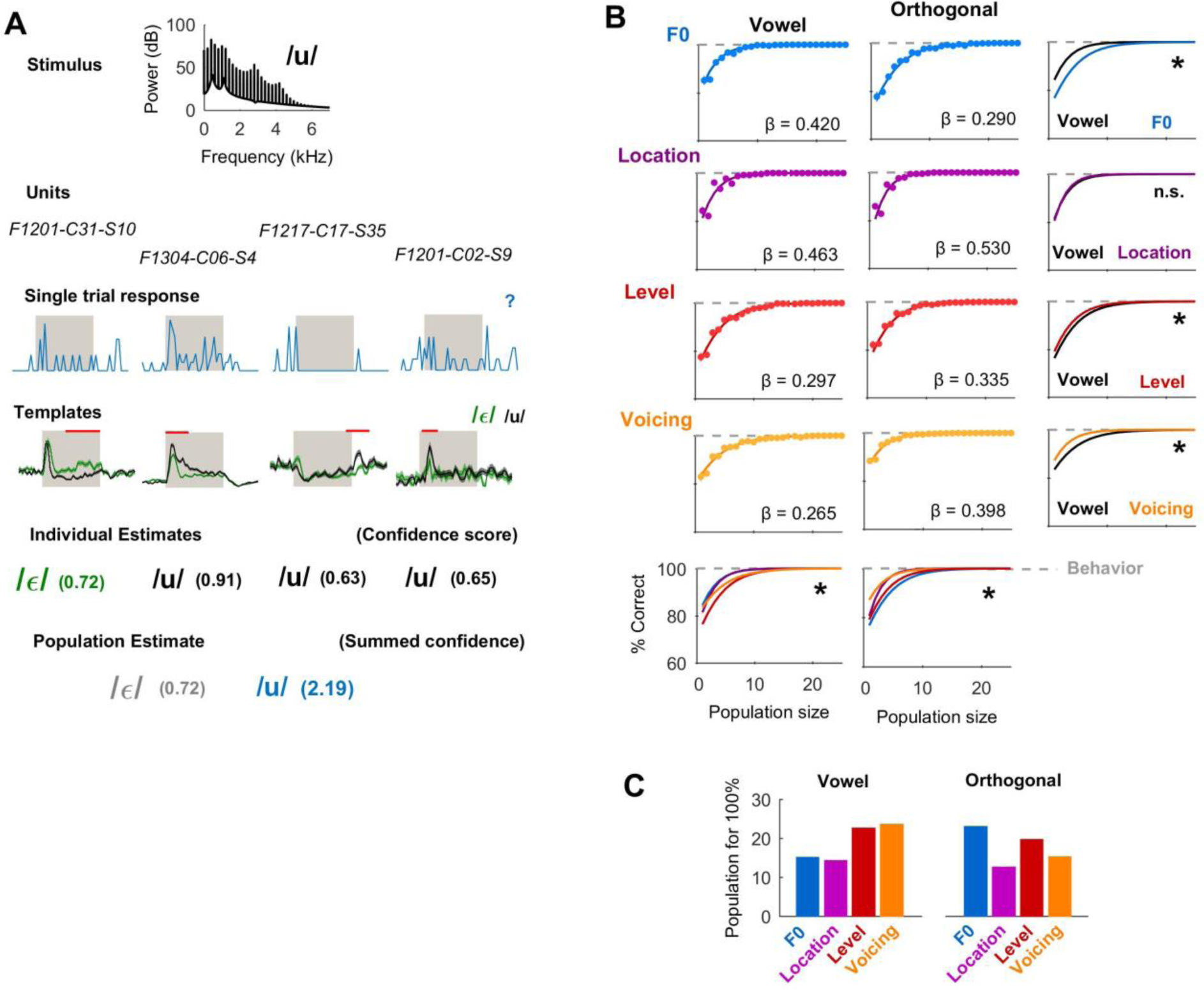
Population decoding can match behavioral performance. **(A)** Schematic illustration of population decoder in which individual unit estimates of acoustic features (e.g. vowel identity) were weighted using a confidence score. Red lines above templates indicate time window of response considered for each unit. **(B)** Decoding performance obtained with increasing population sizes for decoding of vowel identity and orthogonal values as sounds varied in F0, sound location, level or voicing. Data points show mean ± s.e.m. performance of individual populations for each population size. Curves show logistic regression fits with coefficients (β). Asterisks show significant differences (analysis of deviance, *p* < 0.001) between vowel and orthogonal curves (right column) or between orthogonal dimensions (bottom row). Behavioral performance (grey lines) was 100% when considering only correct trials. **(C)** Number of units required to decode variables with performance matching animal behavior.

### Error trials reveal behavioral role for auditory cortex

Population decoding showed that animal’s behavioral performance could be matched by information sampled in the responses of small groups of neurons. We next asked how auditory cortical activity was related to behavior by analysing neural responses on error trials. We reasoned that, if activity was relevant for perception, decoding of sound features should be worse when animals made mistakes; whereas if activity was purely stimulus-driven and independent of behavior, decoding should be similar on correct and error trials, as the same stimuli were presented. Using the timing parameters optimized to decode vowel identity on correct trials, we found that, for vowel-informative units, decoding performance was significantly worse on error than correct trials (Fig 5A-B): this was true for sounds that varied across F0 (Wilcoxon sign-rank: *z* = -8.64, *p* = 5.83 x 10^-18^), location (*z* = 3.57, *p* = 3.61 x 10^-4^), sound level (z = 6.07, *p* = 1.30 x 10^-9^), voicing (*z* = 5.81, *p* = 6.16 x 10^-9^), and when decoding orthogonal values (Fig S6).

The decline in decoding performance we observed on error trials could reflect impairment in the representation of the animals’ choice (respond left/right) rather than vowel identity, as choice and vowel were equivalent on correct trials – as correct trials are defined as those on which a specific vowel produces a specific response (e.g. always respond left to /ε/). If units were purely choice-driven, decoding of the animals’ behavioral response (left or right) should be similar on error and correct trials, while decoding of the stimulus should be significantly worse than chance (50%) as the decoder systematically mis-categorizes trials. While there were many units in which stimulus decoding was substantially below the 50% point, we also saw significantly worse decoding of choice on error trials (Fig 5C-D and Fig S7) when sounds varied across F0 (*z* = -10.1, *p* = 7.50 x 10^-24^), location (*z* = 5.22, *p* = 1.82 x 10^-7^), level (z = 6.93, *p* = 4.19 x 10^-12^) and voicing (*z* = 5.25, *p* = 1.48 x 10^-7^).

To contrast the influence of sensory and choice information on neural activity, we compared the error-related decline in decoding of vowel identity and behavioral choice. Decline in decoding performance was larger for choice than vowel identity when sounds varied across F0 (Fig 5E, rank-sum test, z = 6.54, *p* = 6.09 x 10^-11^), location (z = 2.48, *p* = 0.013) or level (z = 4.11, *p* = 3.89 x 10^-5^) but not voicing (z = 0.473, *p* = 0.636). These findings suggest that, across auditory cortex, neurons provide a predominantly stimulus based representation whose quality determined the animals’ discrimination ability. However, the presence of units in which choice decoding on error trials was maintained, and the observation of units in which decoding of the stimulus on error trials was substantially worse than chance, indicates that the representation is not purely sensory and includes choice information.

**Figure 5.**
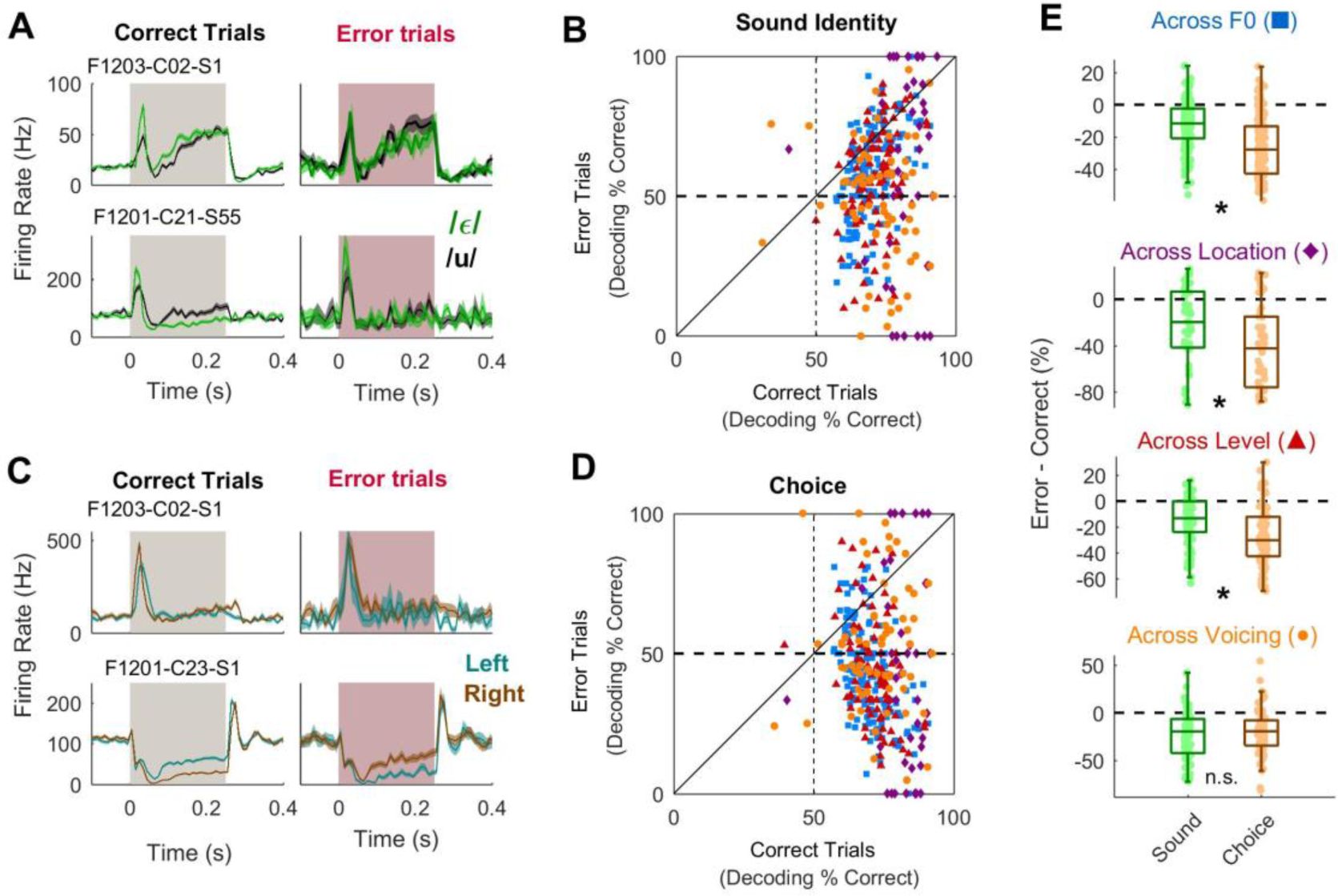
Effects of task accuracy on auditory cortex. (**A**) Discrimination of sound (vowel) identity by individual units on correct and error trials. Bars represent the duration of the first vowel token after stimulus onset; neural responses shown as mean ± s.e.m. firing rates. (**B**) Performance decoding sound identity on correct and error trials for all units. Data presented separately for vowels varied across F0, location, sound level and voicing. (**C**) Discrimination of behavioral choice when animals responded at left or right port by individual units on correct and error trials. Data is shown as in (A). (**D**) Performance decoding behavioral choice on correct and error trials for all units. Data shown as in B. (**E**) Comparison of the effects of task accuracy on decoding sound identity and behavioral choice for vowels varied across each orthogonal dimension. Asterisks show significant differences between sound and choice.

### Choice and accuracy related activity

Our analysis indicated the encoding of both sensory and behavioral variables during perceptual constancy. To study this further, we subsampled neural responses to generate matched datasets containing equal numbers of correct and error trials, vowel identities and choice directions from data across pooled all orthogonal dimensions for which animals showed perceptual constancy (Fig 6A; See Methods). This allowed us to determine modulation of the neural response by the stimulus, the behavioral choice and accuracy (which might indicate confidence or inattention as analysis time windows were restricted to the time before behavioral response). For matched data, behavioral performance would correspond to 50% correct and modulation by one variable (e.g. choice) could not be trivially explained by other variables (e.g. accuracy or sound).

When decoding neural responses from matched data, we confirmed that while information about stimulus identity was more widespread than about behavioral variables, units also conveyed information about choice and accuracy: 37.7% of units (90/239) were significantly informative (permutation test, *p* < 0.05) about sound identity, 23.4% (56/239) about choice and 22.2% (53/239) informative about trial accuracy (Fig 6B). Decoding performance was significantly better for vowel identity than for choice or accuracy (Fig. 6C-D, Kruskal-Wallis test: χ^2^ = 17.5, *p* = 1.58 x 10^-4^; Tukey-Kramer corrected pairwise comparisons: vowel vs. choice, *p* = 0.0024; vowel vs. accuracy, *p* = 3.43 x 10^-4^; choice vs. accuracy, *p* = 0.864). Population decoding plateaued with fewer units when decoding vowel identity (Fig 6E, 18 units required for 100% correct) than decoding accuracy (21 units) or choice (25 units) and performance growth curves differed significantly vowel identity and choice (Bonferroni corrected analysis of deviance, χ^2^ = 154.3, *p* < 0.001), and vowel identity and accuracy (χ^2^ = 109.0, *p* < 0.001). We also compared decoding of choice and accuracy, but found no significant differences (*p* > 0.05) in decoding performance of individual units (Fig 6D) or population decoding functions (Fig 6E). Nonetheless, there were clear representations of behavioral, as well as stimulus, variables, as we could decode the animal’s behavioral choice and accuracy better than chance in a substantial proportion (>20%) of individual units and with perfect decoding performance across small populations.

Our results show that vowel identity was represented across behavioural variables as well as orthogonal stimulus variations, as units represented vowel identity across the animals’ behavioural responses and performance accuracy. Given that auditory cortex multiplexed sound features, we asked if sensory and non-sensory variables were also encoded at different times. We found that information about sound identity emerged earliest, followed by task accuracy and then behavioral choice (Fig 6F): For 147 units that were informative about sound identity, choice and/or accuracy, the time of best decoding differed significantly between dimensions (Kruskal-Wallis test, *χ*^2^ = 15.07, *p* = 5.35 x 10^-4^) with choice represented later than sound identity (Tukey-Kramer corrected, *p* = 3.07 x 10^-4^) but timing of information about accuracy not significantly different from either variable (*p* > 0.1). Thus units multiplexed behavioral, as well as sensory variables, with a sequence consistent with sensory-motor transformation, and provided a tolerant representation of sound identity across orthogonal behavioral, as well as acoustic dimensions.

**Figure 6.**
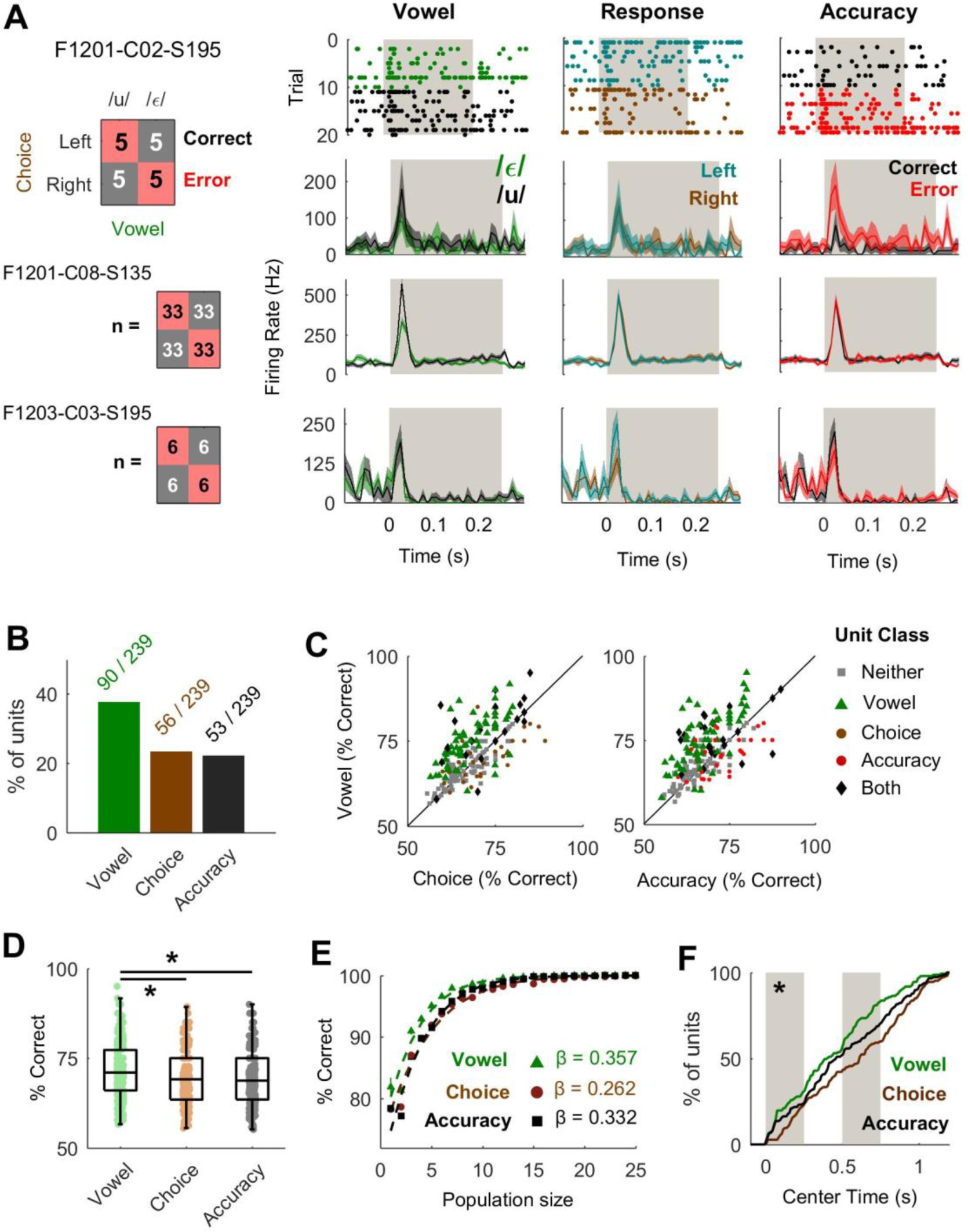
Auditory cortical neurons encode sound identity, behavioral choice and task accuracy. (**A**) Analysis design for matching equal numbers of neural responses to each sound identity, left and right choices, and correct and error trials. Data shown as raster plots of spike times on each trial for one unit and PSTHs representing mean ± s.e.m. firing rate across trials for three units. Grey bars show the first stimulus token. Trial contingency (i.e. respond left for /ε/) shown as an example on which one ferret was trained (F1217). (**B**) Percentage of units informative about sound, choice and/or accuracy in matched data. (**C**) Performance decoding sound, choice and accuracy across all units. (**D**) Comparison of performance decoding sound identity, behavioral choice and task accuracy; boxplots show mean and interquartile range. Lines show significant pairwise comparisons ( *p*< 0.01). (**E**) Performance decoding sound, choice or accuracy with populations of units. Data points show mean ± s.e.m. population performance for each population size. (**G**) Cumulative distributions showing center times for best performance when decoding vowel identity behavioral choice and task accuracy. Grey bars represent the duration of the each token within the stimulus. Data shown for all units informative about one or more variables; asterisk reflects significant difference between variables (*p* < 0.001).

### Effects of Task Engagement

The encoding of animal’s choice and accuracy illustrates that auditory cortex processing extends beyond the representation of acoustic input. Attentional state influences auditory cortical activity (Dong et al., 2013; Kuchibhotla et al., 2017; Otazu et al., 2009) and receptive field properties (Atiani et al., 2014; David et al., 2012; Fritz et al., 2003; Jaramillo and Zador, 2011; Lee and Middlebrooks, 2011; Lu et al., 2017; Niwa et al., 2012). We therefore asked if neural tolerance only emerged during task engagement, by comparing unit responses (e.g. Fig S8) recorded during task performance and during passive listening.

We observed that task engagement suppressed spiking responses in the first 100 ms after stimulus onset (Fig 7A, sign-rank test, z = 3.62, *p* = 2.93 x 10^-4^). In the same time window, we decoded vowel identity significantly better from units recorded task-engaged than passively listening animals (Fig 7B, z = -2.83, *p* = 0.0047). We then expanded our analysis in time to consider effects of engagement with a sliding window, finding that changes in spiking activity and decoding performance were strongly time-dependent: Engagement-related suppression of firing rates occurred throughout stimulus presentation, and contrasted with sustained enhancement of activity in the anticipatory period before stimulus onset (Fig 7C). Furthermore, the *difference in firing rate* between passive and task-engaged units differed significantly with time (one-way anova, F30, 4743 = 7.08, *p* = 1.38 x 10^-28^). Engagement-related enhancement of vowel decoding was observed at the onset and offset of sounds but not in the sustained period of sound presentation (Fig 7D) and the effect of task-engagement varied significantly with time (F30, 4650 = 1.57, *p* = 0.0247).

To understand how the time-dependent effects of task-engagement modulated overall information content, we compared spiking and vowel decoding in the time window that gave best decoding performance, optimised for each condition independently. Consistent with fixed window analyses, firing rates in optimized windows were lower in the engaged than passive condition (Fig 7E; Wilcoxon sign-rank test: z = 3.20, *p* = 0.0014). However, in contrast to findings with fixed time windows, engagement did not improve optimal decoding performance: for units that were significantly vowel informative during active listening, decoding performance was statistically indistinguishable (z = -0.55, p = 0.582, note firing rate difference was still significant for these units, z = 2.41, *p* = 0.016) while if all units were considered, there was a small but significant drop in performance (Fig 7F; z = 2.15, *p* = 0.032). Task engagement similarly affected representation of F0, by supressing spiking activity (z = 2.98, *p* = 0.003) and decoding performance (z = 4.45, *p* = 8.47 x 10^-6^) in optimized time windows (Fig S9).

To understand the origin of differences in fixed-window and optimised analyses, we compared the timing parameters that gave best performance decoding vowel identity, with a focus on units that were significantly vowel informative during task performance. This revealed that the optimized time window for vowel-informative units was significantly earlier during task performance than passive listening (sign-rank test on center time: z = 2.79, *p* = 0.015). The effects of task-engagement were therefore not to enhance the degree of tolerance of vowel informative units, as judged by optimized decoding of vowel identity across F0, which remained similar across states, but rather to enhance the speed and efficiency of tolerant representations, encoding vowel identity faster and using fewer spikes.

**Figure 7.**
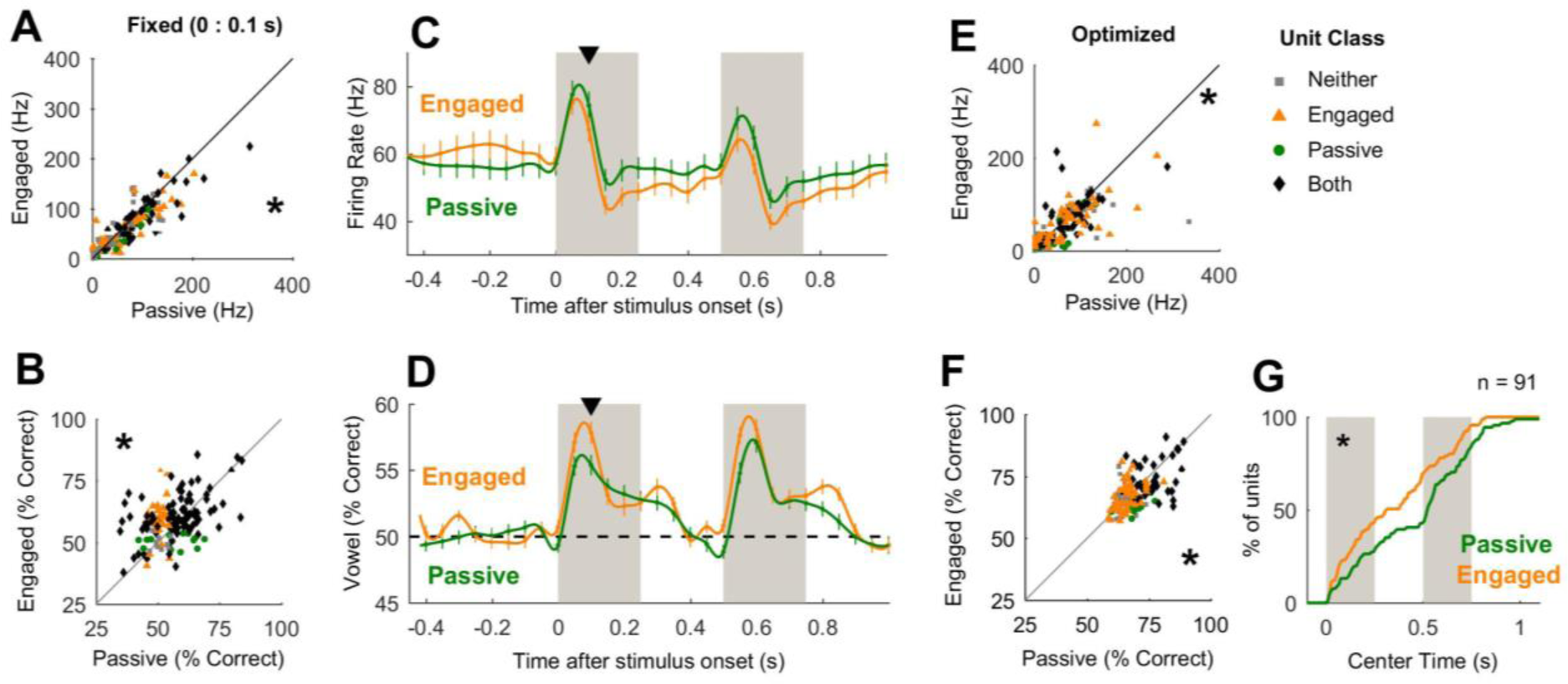
Modulation of auditory processing by task engagement. (**A**) Paired comparison of mean firing rate in the 100 ms after stimulus presentation for units (n = 154) recorded during task performance (engaged) and passive listening conditions. Data points show individual units labelled by classification as informative about vowel identity in engaged and passive conditions. (**B**) Paired comparison of performance decoding vowel identity using neural responses measured in the 100 ms after stimulus onset. Individual data points shown each unit (n = 151). Data is shown as in (A). (**C-D**) Paired comparison of firing rate (C) and vowel decoding performance (D) in time windows fixed relative to stimulus onset. Data points show mean ± s.e.m. Black triangles indicate the comparison at 0 – 100 ms in A-B. (**E**) Firing rate in the time window that gave best performance decoding vowel identity (optimized independently for each unit in each experimental condition [passive/ engaged]). Data is shown as in (A). (**F**) Paired comparison of best performance decoding vowel identity in optimized time windows. Data is shown as in (A). (**G**) Cumulative density distributions showing center times giving best decoding performance. Data is shown for units informative about vowel identity during task performance.

### Effects of Training

In addition to task engagement, long-term experience can also affect auditory cortical processing (Bao et al., 2004; Ohl et al., 2001; Polley et al., 2004; Polley et al., 2006; Schnupp et al., 2006; Whitton et al., 2014)(Atilgan et al. Unpublished) and so we also asked if training to discriminate vowels altered auditory representations. We recorded sound-evoked responses (Fig. S8) to vowels in four naïve ferrets (86 units), and in two trained animals presented with untrained vowels (56 units) and compared these with units recorded in trained animals responding to trained vowels (230 units). As we could not pair units across trained and naïve animals, we conducted unpaired comparisons of neural activity (normalized relative to a pre-stimulus baseline period) and decoding performance.

Training suppressed neural activity in both comparisons of unit responses to trained and untrained stimuli (Fig 8A) and comparisons of units from trained and naïve animals (Fig 8B): Using a roving analysis window and ANOVA to compare normalized firing rates, with time bin and stimulus training as factors, we found significant effects of time (F_30, 8804_ = 29.0, *p* < 0.001), training (F_1, 8804_ = 24.1, *p* < 0.001), and a time x training interaction (F_30, 8804_ = 1.89, *p* = 0.0024). When comparing firing rates in units recorded from trained and naïve subjects, we also found significant effects of time (F_30, 9734_ = 51.3, *p* < 0.001), training (F_1, 9734_ = 25.3, *p* < 0.001), and a time x training interaction (F_30, 9734_ = 3.83, *p* < 0.001).

Training also reduced performance decoding vowel identity across F0 in both comparisons of unit responses to trained and untrained stimuli (Fig 8C), and of units recorded in trained and naïve animals (Fig 8D): Comparisons across time (two-way ANOVA) showed significant effects of stimulus training (F_1, 8804_ = 7.69, *p* = 0.006), time (F_30, 8804_ = 13.6, *p* < 0.001), and a significant time x training interaction (F_30, 8804_ = 1.65, *p* = 0.014). Similarly, subject training (F_1, 9734_ = 12.4, *p* < 0.001) and time (F_30, 9734_ = 16.3, *p* < 0.001) significantly affected vowel decoding – although we found no significant interaction (F_30, 9734_ = 0.73, *p* = 0.857). Thus when assessing the effects of training by comparison of stimuli or subjects, neural activity and vowel decoding performance was suppressed.

Training-related suppression of vowel decoding was also observed when the time parameters of decoding were optimized for each unit (Fig. 8E): Comparing decoding performance across all units recorded in passive conditions revealed a significant effect of experimental group (Kruskal-Wallis test, *χ^2^* = 12.08, *p* = 0.002), with pairwise comparisons revealing significant differences between decoding of responses to trained and untrained sounds (Tukey-Kramer corrected, *p* = 0.046), and between units in trained and naïve animals responding to the same physical stimuli (*p* = 0.007) but not between units in trained and naïve animals responding to unfamiliar sounds (*p* = 0.987). In contrast to fixed time window analysis however, we saw no significant effects of training on firing rates in optimized time windows (Fig 8F, *p* > 0.1). Thus both fixed-window and optimized decoding show training-related reduction in information about vowel identity across F0. Similar results were also found for decoding of F0 across vowels (Fig S10), suggesting that training has broad effects on auditory processing and that information about sound features was, paradoxically, more robust in naïve than trained animals.

**Figure 8.**
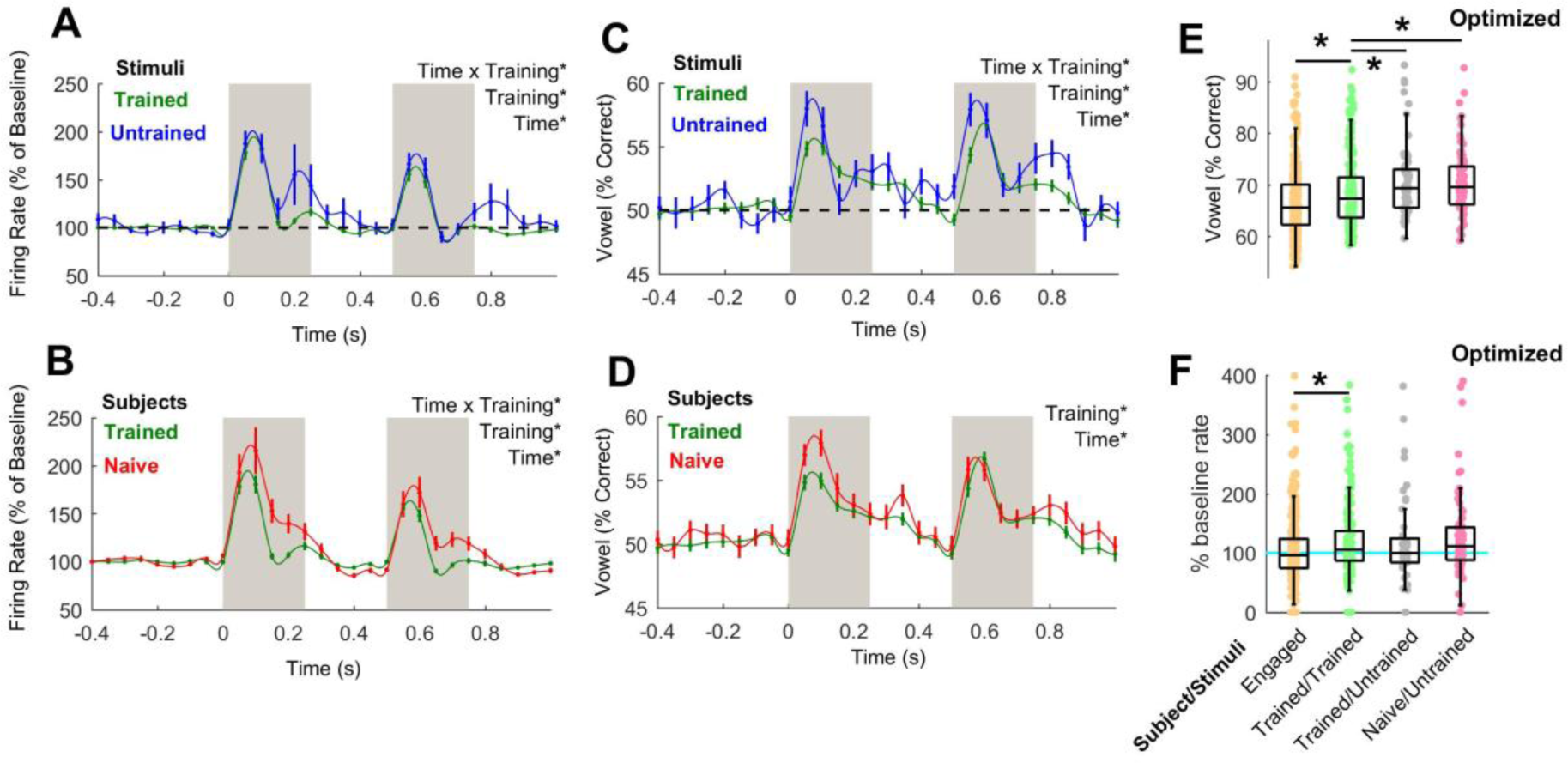
Modulation of auditory processing by training. (**A-B**) Firing rates of units evoked by trained and untrained sounds (A) and in units recorded from trained and naïve animals (B). Data is shown as mean ± s.e.m. in 100 ms windows at 50 ms intervals with spline interpolation across means. (**C-D**) Unpaired comparison of performance decoding vowel identity from unit responses to trained and untrained sounds (C) and units recorded in trained and naïve animals (D). Data is shown as in A-B. (**E**) Comparison of best performance decoding vowel identity in optimized time window. Individual data points show individual units; box plots show median and inter-quartile ranges. Asterisks show significant comparisons between experimental groups (Tukey correction for multiple comparisons, *p* < 0.05). Effect of task engagement shown for reference. (**F**) Normalized firing rate in the time window giving best performance decoding vowel identity. Data is shown as in F.

## Discussion

Here we demonstrate that auditory cortical neurons reliably represent vowel sounds across a range of orthogonal acoustic transformations that mirror those preserved in perceptual constancy. The neural representation provided by auditory cortex was multivariate, as units represented multiple stimulus features, and multiplexed, as variables were best represented at different times. Multivariate encoding extended to behavioral dimensions as units represented subjects’ choice and accuracy and decoding performance differed between correct and error trials. Consistent with a shift from stimulus-related to task-related neural representation, we found that both task-engagement and long-term training significantly affected the representation of vowel identity in auditory cortex. Together our findings demonstrate that auditory cortical neurons provide a degree of tolerance across variation in sensory input and behavior that was sufficient to represent the identity of target sounds during perceptual constancy.

Ferrets identified vowels by their spectral timbre while sounds varied across the major acoustic dimensions key to real-world hearing, including F0 that determines voice pitch, sound location and sound level. Both animals and neurons generalized across the same acoustic dimensions (F0, space etc.). Encoding of multiple features of speech-like sounds, sometimes by the same units, supports previous reports of distributed coding in auditory cortex (Bizley et al., 2009; Griffiths et al., 2010; Ortiz-Rios et al., 2017) and shows that even when potentially disruptive to behavior, orthogonal variables (e.g. F0) are encoded. Furthermore, the encoding of vowel identity by even small populations of units was sufficient to account for, or exceed, animal’s behavioral performance. This suggests that auditory cortex provides a multivariate representation of sounds from which downstream neurons may select behaviorally relevant dimensions during perceptual constancy.

The point at which multivariate encoding of stimulus features might transition to a univariate representation of a task-relevant dimension is unclear. Here we recorded from a combination of primary and secondary tonotopic areas of auditory cortex; however the limited density of electrodes in our recording array prevented us from mapping the precise boundaries between regions necessary to determine if neural tolerance differed between fields as suggested elsewhere (Carruthers et al., 2015). In future it will be important to record using denser arrays with which tolerant representations can be mapped across auditory cortex and beyond: Neurons in prefrontal cortex (PFC) and higher-order auditory cortex (dPEG) are selective for behaviorally relevant sounds (Atiani et al., 2014; Ding and Simon, 2012; Fritz et al., 2010; Russ et al., 2008; Tsunada et al., 2016) and so we might expect that in such areas, tolerant representations of sound identity are preserved while encoding of orthogonal task-irrelevant dimensions is lost.

We decoded vowel identity and orthogonal variables independently and without *a priori* selection of neural response time windows. This approach showed that responses of units informative about both vowel identity and orthogonal features were best decoded in distinct time windows. Temporal multiplexing by units mirrored the time-course of sound perception: Decoding of vowel identity and sound location earlier than voicing or F0 is consistent with perception of sound location and vowel identity at sound onset (Litovsky et al., 1999; Stecker and Hafter, 2002), while listeners require longer to estimate F0 (Gray, 1942; Mckeown and Patterson, 1995; Walker et al., 2011). Best decoding of sound level after vowel identity, sound location or voicing may reflect the time course of temporal integration by the auditory system when assessing moderate level sounds (Buus et al., 1997; Glasberg and Moore, 2002). However, information about voicing was decoded earlier than other stimulus attributes, suggesting that information about harmonicity is available earlier in the neural response, and that temporal multiplexing occurs even when perceptual constancy does not.

The order in which acoustic feature representations emerged during perceptual constancy also matched the encoding of vowel identity, F0 and location under anaesthesia (Walker et al., 2011), indicating that multiplexing is a general principle of encoding in auditory cortex. Our work extends these findings to additional acoustic features (voicing and sound level) as well as non-sensory variables (choice and accuracy). Furthermore, our comparison of engaged and passively listening conditions showed that the time-course of multiplexing was plastic and depended on behavioral state. By accelerating the encoding of acoustic variables during task performance, neurons may create time for integration of motor and motivational signals, as well as taught associations (Fritz et al., 2003; Fritz et al., 2010; McGinley et al., 2015; Schneider et al., 2014) in order to coordinate behavioral responses. We would therefore predict that delaying the encoding of acoustic features, but preserving the overall information content of auditory cortical responses, would either disrupt or retard sound discrimination.

An open question is why training animals to discriminate sounds reduced information about stimulus features. Such effects are consistent with independent findings that training animals to discriminate vowel identity leads to a reduction in the variation in auditory cortical responses attributable to vowel identity and F0 (Atilgan et al., Unpublished). Furthermore, changes in decoding performance could not be explained trivially by changes in firing rate, as we observed both suppression of neural activity and enhancement of decoding performance (Fig 7), and suppression of decoding performance in the absence of changes in neural activity (Fig 8). One possibility could be that responses to untrained sounds reflect purely feedforward information about sound features extracted earlier in the auditory pathway, but that the association of sounds with non-sensory dimensions in auditory cortex comes at the cost of representing acoustic information.

Our findings confirm the importance of behavioral variables in auditory cortical processing (Bizley et al., 2013b; Dong et al., 2013; Niwa et al., 2012): decoding of sound features was impaired on error trials, and we found many units that encoded information about the animals’ choice and /or accuracy. The significant drop in decoding performance on error trials, and the sensitivity of units to accuracy, shows that auditory cortical activity is predictive of upcoming mistakes. Given this information, and the finding that stimulus identity could be decoded perfectly from small populations of units, why do animals make errors? One possibility is that errors arise from inattention, which has a distinct neural signature (Lakatos et al., 2016) that our decoder uses to distinguish correct and error trials. At present it is unclear whether the accuracy signal we decode reflects such an attentional lapse or arises as an interaction between representations of sound identity and behavioral choice, or a representation of confidence in auditory processing, or anticipation of reward (Metzger et al., 2006). Future experiments in which confidence or reward value are systematically explored may explain the precise nature of accuracy information reported here.

In summary, our results show that during perceptual constancy, neurons in auditory cortex provide tolerant representations of vowel identity and that small populations of units can represent sounds as well as, or better than animal’s behavior. Auditory cortical units also encoded information about F0, sound location, level and voicing, as well as the animal’s choice and accuracy in the task, each with a specific temporal profile that shows a multivariate and multiplexed system. Task-engagement and training modulated auditory processing, demonstrating a role for attention and long-term experience in perceptual constancy. Across all these variables and experimental conditions, auditory cortical responses showed sufficient tolerance to unambiguously represent vowel identity in the same conditions that animals successfully generalized behavioral performance, and thus provided a neural correlate of perceptual constancy.

## Author contributions

SMT and JKB designed the experiments and wrote the paper; all authors were involved in data collection; SMT analysed the data.

## Acknowledgements

This work was funded by a Royal Society Dorothy Hodgkin Fellowship to JKB, the BBSRC (BB/H016813/1) and the Wellcome Trust / Royal Society (WT098418MA).

## Methods

### Animals

Subjects were four pigmented female ferrets (1-5 years old) trained to discriminate vowels across fundamental frequency, sound level, voicing and location (Bizley et al., 2013a; Town et al., 2015). Each ferret was chronically implanted with Warp-16 microdrives (Neuralynx, MT) housing sixteen independently moveable tungsten microelectrodes (WPI Inc., FL) positioned over primary and posterior fields of left and right auditory cortex (Fig S2). Details of the surgical implantation procedures and histological confirmation of electrode position are described elsewhere (Bizley et al., 2013c). A further six ferrets (also pigmented females) implanted with the same microdrives were used as naïve animals for passive recording. These animals were trained in a variety of psychophysical tasks that did not involve the vowel sounds presented here.

Subjects were water restricted prior to testing; on each day of testing, subjects received a minimum of 60ml/kg of water either during testing or supplemented as a wet mash made from water and ground high-protein pellets. Subjects were tested in morning and afternoon sessions on each day for up to five days in a week. Test sessions lasted between 10 and 50 minutes and ended when the animal lost interest in performing the task.

The weight and water consumption of all animals was measured throughout the experiment. Regular otoscopic examinations were made to ensure the cleanliness and health of ferrets’ ears. Animals were housed in groups of two or more animals in enriched housing conditions. All experimental procedures were approved by a local ethical review committee and performed under license from the UK Home Office and in accordance with the Animals (Scientific Procedures) Act 1986.

### Apparatus

Ferrets were trained to discriminate sounds in a customized pet cage (80 cm x 48 cm x 60 cm, length x width x height) within a sound-attenuating chamber (IAC) lined with sound-attenuating foam. The floor of the cage was made from plastic, with an additional plastic skirting into which three spouts (center, left and right) were inserted. Each spout contained an infra-red sensor (OB710, TT electronics, UK) that detected nose-pokes and an open-ended tube through which water could be delivered.

Sound stimuli were presented through two loud speakers (Visaton FRS 8) positioned on the left and right sides of the head at equal distance and approximate head height. These speakers produce a smooth response (±2 dB) from 200Hz to 20 kHz, with an uncorrected 20 dB drop-off from 200 to 20 Hz when measured in an anechoic environment using a microphone positioned at a height and distance equivalent to that of the ferrets in the testing chamber. An LED was also mounted above the center spout and flashed (flash rate: 3 Hz) to indicate the availability of a trial. The LED was continually illuminated whenever the animal successfully made contact with the IR sensor within the center spout until a trial was initiated. The LED remained inactive during the trial to indicate the expectation of a peripheral response and was also inactive during a time-out following an incorrect response.

The behavioral task, data acquisition, and stimulus generation were all automated using custom software running on personal computers, which communicated with TDT real-time signal processors (RZ2 and RZ6, Tucker-Davis Technologies, Alachua, FL).

### Task Design, Stimuli and Behavioral Testing

Ferrets discriminated vowel identity in a two-alternative forced choice task described elsewhere (Town et al., 2015). Briefly, on each trial the animal was required to approach the center spout and hold head position for a variable period (0 – 500 ms) before stimulus presentation. Each stimulus consisted of a 250 ms artificial vowel sound repeated once with an interval of 250 ms. Animals were required to maintain contact with the center spout until the end of the interval between repeats (i.e. 500 – 1000 ms after initial nose-poke) and could then respond at either left or right spout. Correct responses were rewarded with water delivery whereas incorrect responses led to a variable length time-out (3 - 8 s). To prevent animals from developing biases, incorrect responses were also followed by a correction trial on which animals were presented with the same stimuli. Correction trials and trials on which the animal failed to respond within the trial window (60 s) were not analysed. The only exception to this protocol was for whispered sounds, which we presented as probe sounds in 10 – 20% of trials on which any response was rewarded and correction trials did not follow.

We also tested subjects under passive listening conditions, in which animals were provided with water at the center port to recreate the head position and motivational context occurring during task performance. Sounds were presented with the same two-token stimulus structure as during task performance, with a minimum of 1 second between stimuli. During test sessions, sound presentation began once the animal approached the center spout and began licking and ended when the animal became sated and lost interest in remaining at the spout.

Stimuli were artificial vowel sounds synthesized in MATLAB (MathWorks, USA) based on an algorithm adapted from Malcolm Slaney’s Auditory Toolbox (https://engineering.purdue.edu/∼malcolm/interval/1998-010/). The adapted algorithm simulates vowels by passing a sound source (either a click train, to mimic a glottal pulse train for voiced stimuli, or broadband noise for whispered stimuli) through a biquad filter with appropriate numerators such that formants are introduced in parallel. Four formants (F1-4) were modelled: three subjects were trained to discriminate /u/ (F1-4: 460, 1105, 2857, 4205 Hz) from /ε/ (730, 2058, 2857, 4205 Hz) while one subject was trained to discriminate /a/ (936, 1551, 2975, 4263 Hz) from /i/ (437, 2761, 2975, 4263 Hz). Selection of formant frequencies was based on previously published data (Peterson and Barney, 1952; Town et al., 2015) and synthesis produced sounds consistent with the intended phonetic identity. Formant bandwidths were kept constant at 80, 70, 160 and 300 Hz (F1-4 respectively) and all sounds were ramped on and off with 5 ms cosine ramps.

To test perceptual constancy, we varied the rate of the pulse train to generate different fundamental frequencies and used broadband noise rather than pulse train to generate whispered vowel. For sound level we simply attenuated signals in software prior to stimulus generation. For sound location, we presented vowels only from the left or right speaker whereas for all other tests sounds were presented from both speakers. Across variations in F0, voicing and space, we fixed sound level at 70 dB SPL. For tests across sound level and location, voiced vowels were generated with 200 Hz fundamental frequency. Sound levels were calibrated using a Brüel & Kjær (Norcross, USA) sound level meter and free-field [1/2] inch microphone (4191) placed at the position of the animal’s head during trial initiation.

### Neural Recording

Neural activity in auditory cortex was recorded continuously throughout task performance. On each electrode, voltage traces were recorded using TDT System III hardware (RX8 and RZ2) and OpenEx software (Tucker-Davis Technologies, Alachua, FL) with a sample rate of 25 kHz. For extraction of action potentials, data were bandpass filtered between 300 and 5000 Hz and motion artefacts were removed using a decorrelation procedure applied to all voltage traces recorded from the same microdrive in a given session (Musial et al., 2002). For each channel within the array, we identified spikes (putative action potentials) as those with amplitudes between -2.5 and -6 times the RMS value of the voltage trace and defined waveforms of events using a 32-sample window centered on threshold crossings.

In the current study, waveform shapes were not sorted and data from multiple test sessions combined across days. The activity for each unit thus represents the unsorted multi-unit activity of a small population of cells at the recording site. We identified sound responsive units in task-engaged animals as those whose stimulus evoked response within the 300 ms after onset of first token differed significantly from spontaneous activity in the 300 ms before making contact with the spout (Sign-rank test, *p* < 0.05). In passive conditions, we identified responsive units using a similar comparison, but spontaneous activity was measured in the 300 ms before stimulus presentation.

### Decoding procedure

We decoded stimulus features (e.g. vowel identity, F0 etc.) on single trials using a simple spike-distance decoder with leave-one-out cross-validation (LOCV). For every trial over which an individual unit was tested in a given dataset (e.g. vowels varied across F0 during task performance), we calculated template responses for each stimulus class (e.g. each vowel or each F0) as the mean PSTH of responses on all other trials. We then estimated the stimulus feature on the test trial as the template with the smallest Euclidean distance to the test trial (Fig S1A). Where equal distances were observed between test trial and multiple templates, we randomly estimated (i.e. guessed) which of the equidistant templates was the true stimulus feature. This procedure was repeated for all trials and decoding performance was measured as the percentage of trials on which the stimulus feature was correctly recovered. Although this approach was simple and did not account for the variance of neural activity, it provided a simple and intuitive relationship between neural activity and information content that we could use with small data sets (sample sizes down to five trials per condition). Robustness to sample size was particularly important because the animal’s behavior determined the number of trials in each condition and we aimed to analyse as many units as possible rather than develop a more sophisticated decoder.

Auditory cortical units showed a wide variety of response profiles that made it difficult to select a single fixed time window over which to decode neural activity. To accommodate the heterogeneity of auditory cortical neurons and identify the time at which stimulus information arose, we repeated our decoding procedure using different time windows (n = 1550) varying in start time (-0.5 to 1 s after stimulus onset, varied at 0.1 s intervals) and duration (10 to 500 ms, 10 ms intervals) (Fig S1B). Within this parameter space we then reported the parameters that gave best decoding performance. Where several parameters gave best performance we reported the time window with earliest start time and shortest duration.

To assess the significance of decoding performance, we conducted a permutation test in which the decoding procedure (including temporal optimization) was repeated 100 times but with vowel identity randomly shuffled between trials to give a null distribution of decoder performance (Fig S1C). The null distribution of shuffled decoding performance was then parameterized and fitting a Gaussian probability density function, for which we then calculated the probability of observing the real decoding performance. Units were identified as informative when the probability of observing the real performance was less than 0.05. Parameterization of the null distribution was used to reduce the number of shuffled iterations over which decoding was repeated. This was necessary because the optimization search for best timing parameters dramatically increased the computational demands of decoding.

### Population Decoding

To decode vowel identity from the single trial responses of populations of units, we simply the summed the number of units that estimated each stimulus, weighted by the confidence of each unit’s estimate, and took the stimulus with the maximum value as the population estimate. Weights for individual unit (w) estimates were calculated as

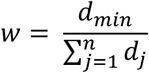

Where *n* was the number of stimulus classes (e.g. vowel identities) and *d* was the spike distance between a test trial response and response templates generated for each stimulus class. Here, d_min_ represents the minimum spike distance that indicated the estimated stimulus for that unit.

We tested populations of up to 35 units, by which point decoder performance had typically saturated at 100% (with the exception of decoding F0 and sound level across larger [n = 5] numbers of feature classes [e.g. 149, 200, 263, 330 and 459 Hz]). Populations were constructed first by selecting the top 35 units that performed best at decoding the relevant parameter at the individual unit level. Within this subpopulation, we randomly sampled 100 combinations of units without replacement from the large number of possible combinations of units available.

### Data Analysis

#### Behavior

Perceptual constancy was reported when the orthogonal factor did not significantly affect task performance, i.e. the likelihood of responding correctly. To test this, we analysed the proportion of correct trials as a function of each orthogonal dimension (e.g. F0) using a logistic regression (Table S1). Regressions were performed separately for each animal, and each orthogonal dimension, and any significant effect (p < 0.05) was reported as a failure of constancy. We also asked if an animal’s performance at specific orthogonal values was better than chance (50%) using a binomial test (p < 0.001, Table S2).

#### Neural activity

The times of spikes was referenced to the onset of the stimulus on each trial and used to create raster and peri-stimulus time histograms. In our analysis of task engagement and training, we measured on each trial the firing rate in 100 ms bins after stimulus onset at 50 ms intervals. For paired comparisons, firing rates in engaged and passively listening animals was compared using a Wilcoxon sign-rank test. For unpaired analyses, we normalized firing rates in these bins relative to the firing rate in a pre-stimulus baseline period in the 450 ms before stimulus onset (passively listening animals) or before the animal began waiting at the center spout (task-engaged animals). Across passively listening groups presented with familiar / unfamiliar sounds (Fig 8E), we compared normalized firing rates and baseline firing rates (i.e. the normalization factors in each condition) across groups using a Kruskal-Wallis test with pairwise post-hoc comparisons performed with Tukey-Kramer correction for multiple comparisons.

#### Individual unit decoding

In addition to classifying whether units were informative about a particular stimulus feature (permutation test, *p* < 0.05), we also compared decoding performances (Fig 5B, 5D, 6D, 7B, 7F, 8E, S6E, S7, S9A and S10A). When comparing decoding performance across more than two conditions (i.e. in passively listening animals; Fig 8E), data were analysed using a Kruskal-Wallis test with Tukey-Kramer corrected post-hoc comparisons where relevant. When comparing two conditions directly, we used a Wilcoxon sign-rank test for paired data (e.g. comparing performance on correct and error trials; Fig 5B). For comparison of changes in decoding performance between conditions (e.g. decoding sound identity vs. choice on correct and error trials; Fig. 5E), we used a Wilcoxon rank-sum comparison for unpaired data.

#### Timing

For each unit, we determined the timing window for which we achieved best decoding performance (Fig S1B) and took the window center (Fig 3), start time (Fig S4) or window duration (Fig S5). We then compared the change in parameter value (e.g. change in center time) for best decoding of between vowel identity and orthogonal dimensions using a Wilcoxon rank-sum test (Fig 3A-D). The same approach was used when comparing the timing of decoding vowel identity and F0 in task-engaged and passively listening animals (Fig 7G). We also compared the times of best decoding of vowel identity across orthogonal dimensions using a Kruskal-Wallis anova with Tukey-Kramer correction for post-hoc comparisons (Fig 3E). We used the same approach to compare the decoding of orthogonal dimensions (Fig 3F) and decoding of vowel identity, behavioral choice and accuracy (Fig 6F).

#### Population decoding

For each unit in a given population, we generated estimates of the target value on each trial based on the minimum spike distance from templates generated on all other trials (i.e. the same LOCV method as for individual unit decoding – see above). Templates were generated using the timing parameters that gave best decoding in the individual unit case and thus each unit’s response was sampled independently. In addition to an estimated target value, we also retained a confidence score for that estimate: the spike distance from test trial to the closest template, expressed as a proportion of the sum of spike distances between test trial and all templates. Across the population, we then summed confidence scores for each possible target value and selected the value with the largest sum as the population estimate for a given trial. We then repeated the procedure across trials to get the decoding performance of a given population.

We summarized the relationship between population size and decoding performance by fitting a logistic regression model to the proportion each population scored correct, with population size as a predictor. To compare population decoding across conditions (e.g. decoding vowel identity or sound location; Fig 4B) we fitted a logistic model with and without the condition as an additional predictor and assessed significance of improvement in model fit using an analysis of deviance.

#### Error trial analysis

We trained the decoder on correct trials using the LOCV procedure to estimate vowel identity on each individual correct trial from templates built on all-other correct trials. For error trials, we used the training templates calculated across all correct trials and estimated vowel identity on each error trial. Only units that were informative about vowel identity were analysed, with the exception of three units recorded when the animal performed perfectly (i.e. made no errors) when vowels varied across sound location and thus error trials could not be studied. We repeated the same procedure for decoding orthogonal variables using only units informative about the relevant dimension. Decoding performance was compared for vowel identity (Fig 5B), orthogonal values (Fig S6) and for behavioral choice (Fig. 5D) using a Wilcoxon sign-rank test. We compared the change in decoding performance between correct and error trials when decoding vowel identity and behavioral choice using a Wilcoxon rank-sum test (Fig 5E).

#### Datasets matched for vowel, choice and accuracy

To study the tolerance of a given unit to behavioral as well as acoustic variables, we subsampled neural responses from all conditions in which animals showed perceptual constancy: Specifically we included sounds varied across F0, sound location and sound level above 60 (three ferrets) or 70 dB SPL (one ferret). We excluded all data when sounds were whispered. To prevent trial outcome (water reward or timeout) from confounding accuracy signals, we also excluded trials on which animals responded within one second of stimulus onset. Following pooling and exclusion, we balanced data sets for the number of each vowel, choice and trial outcome by randomly selecting *N* trials, where N was the minimum number of trials in which any one condition (e.g. left responses to /u/) was tested. As with our earlier decoding analysis, we only considered units for which N ≥ 5. We then decoded vowel identity, behavioral choice and accuracy using the same LOCV decoding procedure described above. We compared decoding performance for vowel identity, choice and accuracy across all units with a Kruskal-Wallis anova and post-hoc comparisons using the Tukey-Kramer correction (Fig 6D).

## Supplemental Tables

**Table S1.** Results of logistic regressions comparing performance across orthogonal variables.

| Ferret | Orthogonal Dimension |  |  |  |
| --- | --- | --- | --- | --- |
|  | F0<br>(149, 200, 263, 330 &<br>459 Hz) | Voicing<br>(Voiced /<br>Whispered) | Sound Level<br>(45 – 82.5 dB SPL) | Location<br>(±90°)<br>(i.e. Left / Right) |
| F1201 | *df* = 12034, *p* = 0.7731 | *df* = 5802, *p* < 0.001 | *df* = 3490, *p* < 0.001 | *df* = 879, *p* = 0.219 |
| F1203 | *df* = 9945, *p* = 0.764 | *df* = 4212, *p* < 0.001 | *df* = 3063, *p* = 0.005 | *df* = 701, *p* = 0.185 |
| F1217 | *df* = 4790, *p* = 0.368 | *df* = 3784, *p* < 0.001 | *df* = 2214, *p* < 0.001 | *df* = 145, *p* = 0.523 |
| F1304 | *df* = 1485, *p* = 0.388 | *df* = 617, *p* = 0.002 | *df* = 455, *p* = 0.882 | *Not tested* |

**Table S2.** Comparison of observed vowel discrimination against chance performance (50%); data shown as fraction of trials correct and probability of observed performance (binomial test). Orthogonal values tested separately for voicing and sound level when a significant main effect of orthogonal value was observed on behavioral performance (Table S1).

| Dimension | Data Set | Ferret |  |  |  |
| --- | --- | --- | --- | --- | --- |
|  |  | F1201 | F1203 | F1217 | F1304 |
| F0 | All | 10214 / 12036<br>84.9%<br>$p < 0.001$ | 8502 / 9947<br>85.5%<br>$p < 0.001$ | 3946 / 4792<br>82.4%<br>$p < 0.001$ | 1051 / 1487<br>70.7%<br>$p < 0.001$ |
| Voicing | Voiced | 3830 / 4472<br>85.6%<br>$p < 0.001$ | 2833 / 3207<br>88.3%<br>$p < 0.001$ | 2585 / 3038<br>85.1%<br>$p < 0.001$ | 333 / 473<br>70.4%<br>$p < 0.001$ |
| | Whispered | 870 / 1332<br>65.3%<br>$p < 0.001$ | 596 / 1007<br>59.2%<br>$p < 0.001$ | 386 / 748<br>51.6%<br>$p = 0.400$ | 83 / 146<br>56.9%<br>$p = 0.116$ |
| Sound level<br>(dB SPL) | 45 | 138 / 186<br>74.2%<br>$p < 0.001$ | 149 / 196<br>76.0%<br>$p < 0.001$ | Not tested | Not tested |
| | 52.5 | 151 / 193<br>78.2%<br>$p < 0.001$ | 169 / 205<br>82.4%<br>$p < 0.001$ | Not tested | Not tested |
| | 60 | 161 / 192<br>83.9%<br>$p < 0.001$ | 162 / 188<br>86.2%<br>$p < 0.001$ | Not tested | Not tested |
| | 64.5 | 438 / 498<br>88.0%<br>$p < 0.001$ | 372 / 414<br>89.9%<br>$p < 0.001$ | 341 / 442<br>77.2%<br>$p < 0.001$ | Not tested |
| | 67.5 | 175 / 195<br>89.7%<br>$p < 0.001$ | 176 / 198<br>88.9%<br>$p < 0.001$ | Not tested | Not tested |
| | 69 | 479 / 523<br>91.6%<br>$p < 0.001$ | 357 / 397<br>89.9%<br>$p < 0.001$ | 345 / 449<br>76.8%<br>$p < 0.001$ | Not tested |
| | 73.5 | 460 / 515<br>89.3%<br>$p < 0.001$ | 352 / 407<br>86.5%<br>$p < 0.001$ | 380 / 457<br>83.2%<br>$p < 0.001$ | Not tested |
| | 75 | 168 / 187<br>89.8%<br>$p < 0.001$ | 179 / 197<br>90.9%<br>$p < 0.001$ | Not tested | Not tested |
| | 78 | 445 / 501<br>88.8%<br>$p < 0.001$ | 382 / 434<br>88.0%<br>$p < 0.001$ | 392 / 442<br>88.7%<br>$p < 0.001$ | Not tested |
| | 82.5 | 445 / 500<br>89.0%<br>$p < 0.001$ | 357 / 424<br>84.2%<br>$p < 0.001$ | 366 / 424<br>86.3%<br>$p < 0.001$ | Not tested |
| | All | Not tested | Not tested | Not tested | 309 / 455<br>67.9%<br>$p < 0.001$ |
| Location | All | 701 / 879<br>79.8%<br>$p < 0.001$ | 564 / 703<br>80.2%<br>$p < 0.001$ | 112 / 145<br>77.2%<br>$p < 0.001$ | Not tested |

## Supplemental Figures

**Fig S1.**
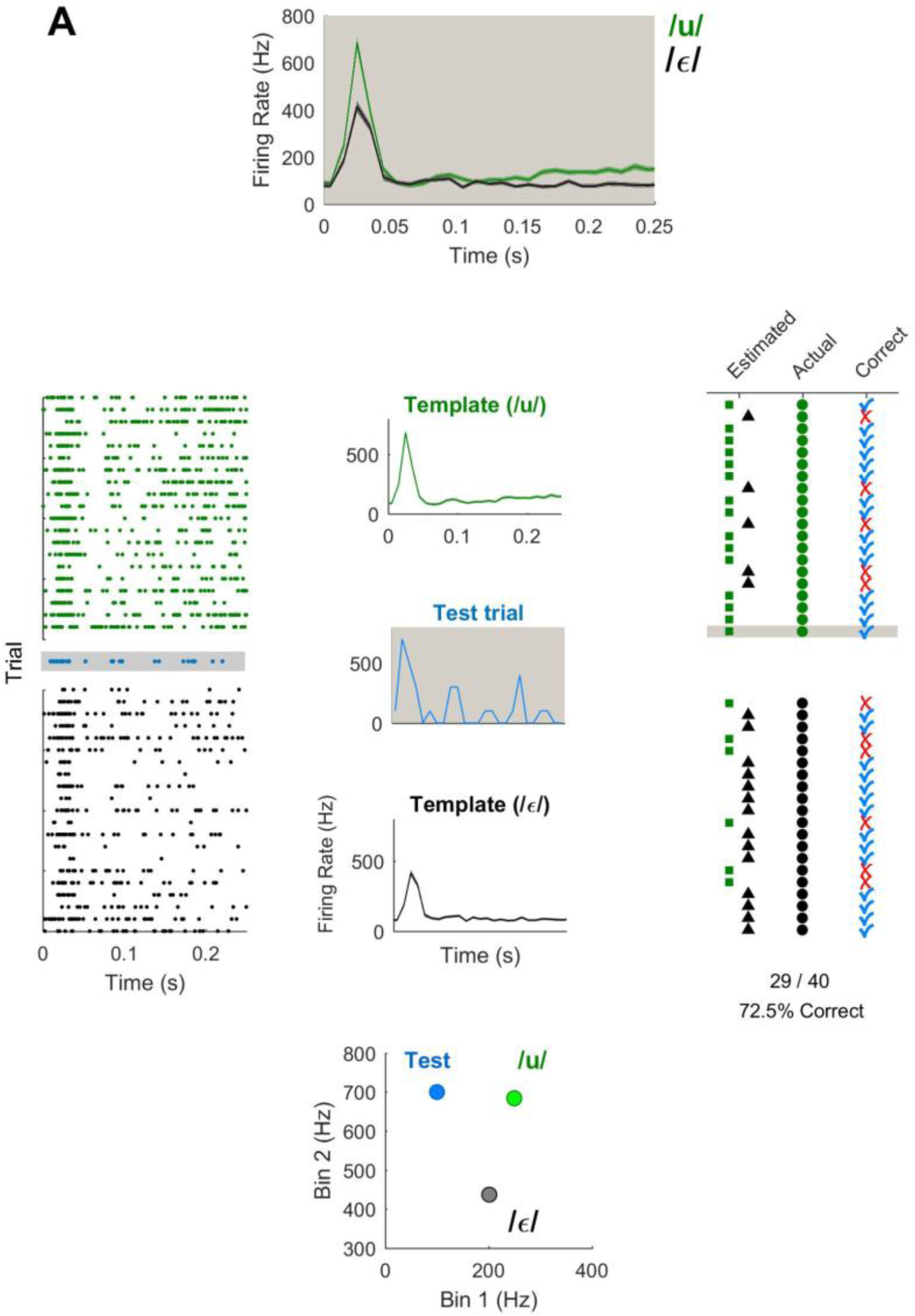
Decoder structure. **(A)** Schematic showing the decoding of trial parameters (e.g. vowel identity) from single trial neural responses for one unit. We used a leave-one-out cross validation method in which templates were calculated as the mean response to each stimulus class (e.g. vowel) on all but one test trial of the data set. Mean responses were averaged across trials from spike times within a decoding window binned at 10 ms intervals. For the test trial, the decoded estimate of stimulus class was assigned as the template class with the smallest Euclidean distance to the test response. Every trial in the dataset was decoded as a test trial with templates recalculated from all other trials.

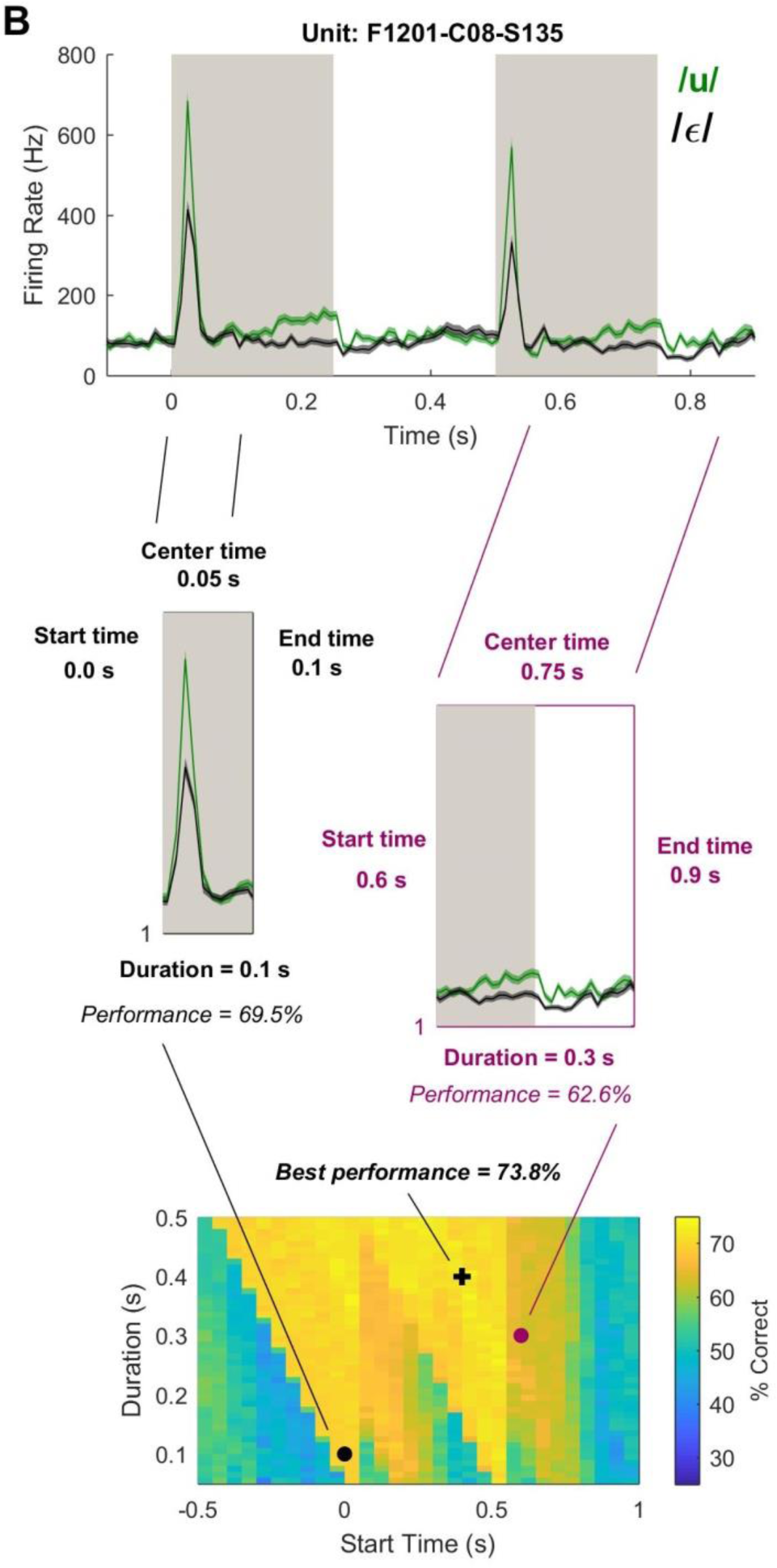
**(B)** Optimizing timing parameters. To accommodate potential variation in timing of information content, we varied the temporal parameters (start time and duration) that defined the decoding window. Start time was varied from -0.5 to 1 s after stimulus onset in 50 ms intervals; duration was varied between 10 and 500 ms in 10 ms intervals. For each combination of start time and duration, we calculated decoding performance across trials and mapped temporal parameter space using a simple grid search. While this search protocol may not find the true optimal parameters for best decoding performance, it nonetheless enabled us to improve decoding performance and estimate those times in the trial at which information about a given feature was most strongly represented.

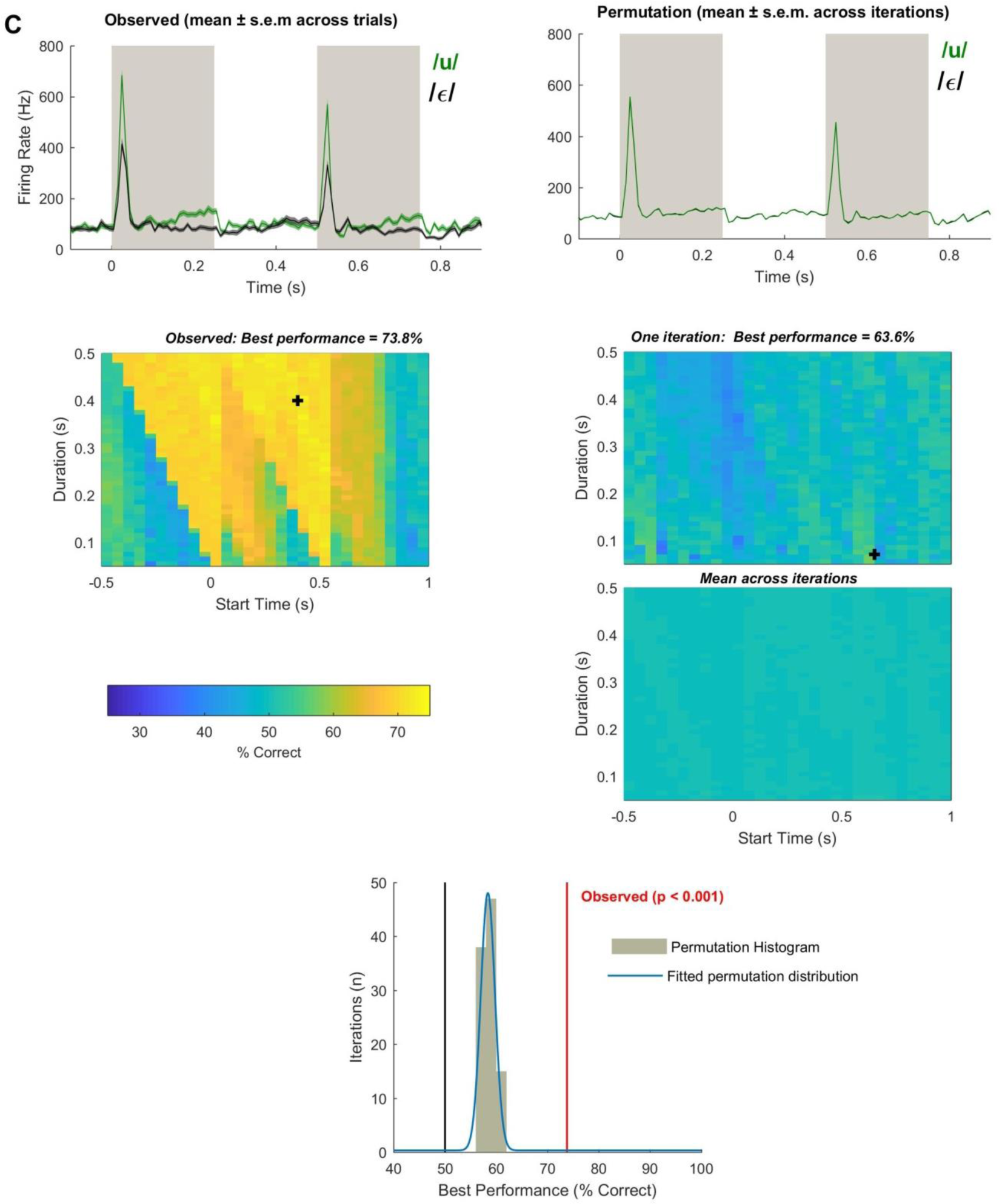
**(C)** Permutation testing of decoder performance. Each test variable (e.g. vowel identity) was shuffled and the decoding was repeated with the full optimization search. For each unit, we repeated this shuffling procedure on 100 iterations (we used a relatively small number of iterations and parameterized the permutation distribution to compromise for the computational cost of optimization). When shuffled, PSTH responses to each vowel were virtually identical. To determine whether a unit was informative, we fitted the distribution of best performance values obtained for each shuffle and calculated the probability of measuring the observed decoding performance.

**Fig S2.**
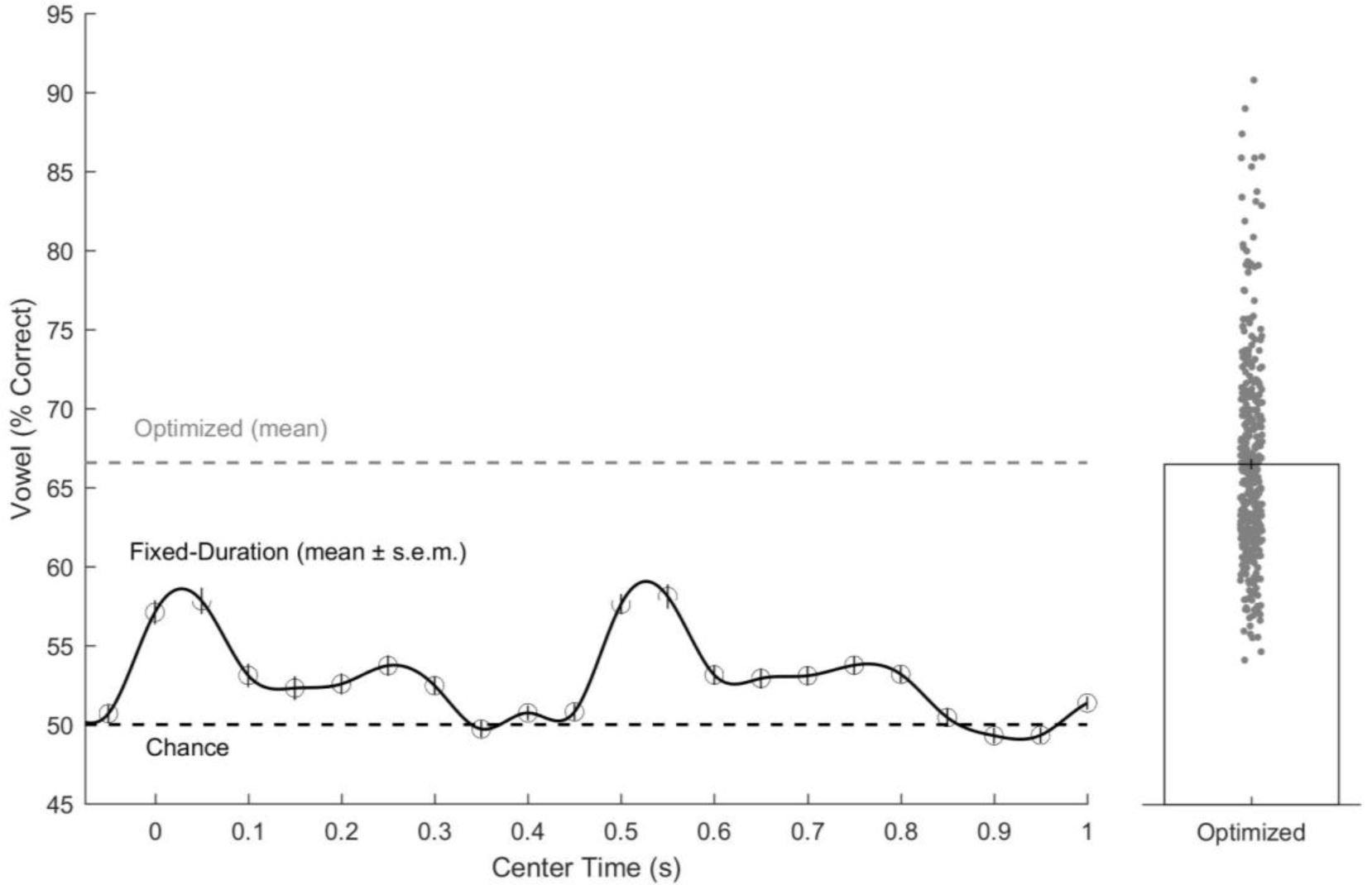
Improvement in decoding performance with optimization of time window parameters. Comparison of performance decoding vowel identity using neural responses in a fixed duration (100 ms centered at different times after stimulus onset) or using optimized time window. Data show mean ± s.e.m. with individual data points showing individual units for optimized data. For each time point, optimized decoding performance was significantly better than fixed window performance (Bonferroni corrected for multiple comparisons, *p* < 1 x 10^-10^).

**Fig S3.**
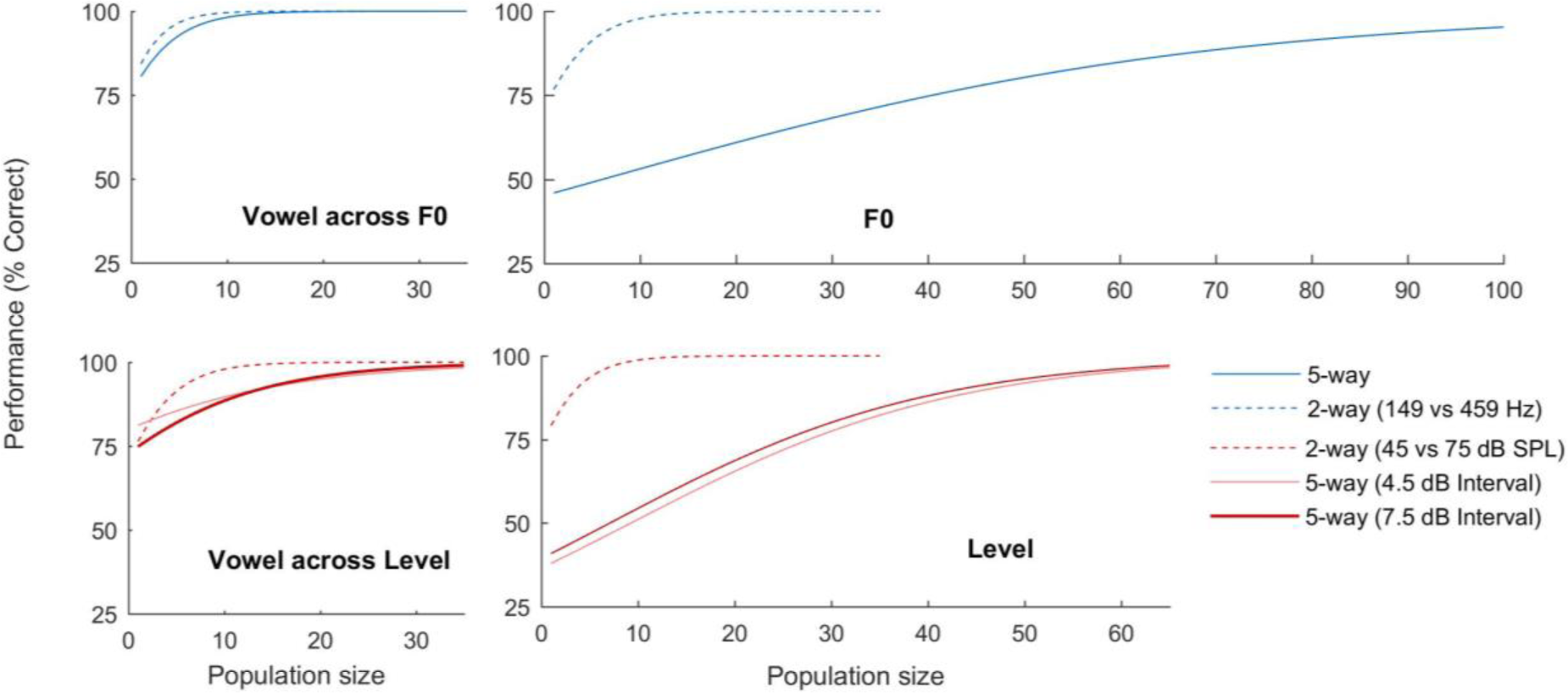
Decoder performance vs. feature set size. Population decoding performance for two and five way classification. Data shown as logistic regression model fits for all populations tested (see Fig 5 of the main text for examples of original data).

**Fig S4.**
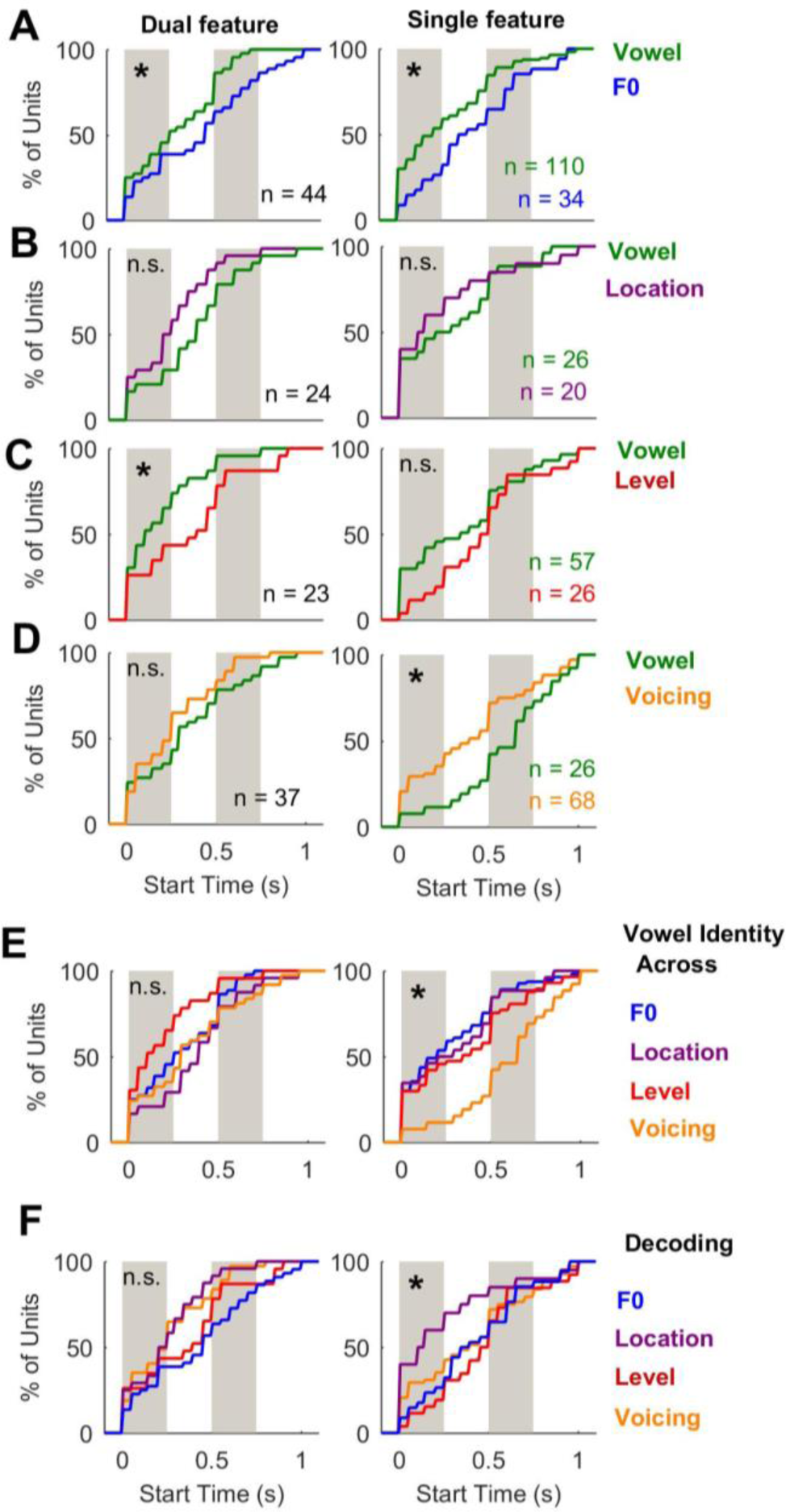
Temporal profiles for the onset (start time) of best decoding windows. **(A-D)** Cumulative distributions showing start time for best performance when decoding vowel identity or orthogonal variables (**A:** F0, **B**: location, **C**: level and **D**: voicing). Units are shown separately by classification as informative about vowel identity and orthogonal values (Dual feature units), or only vowel identity or orthogonal values (Single feature units). **(E)** CDFs for decoding vowel identity across each orthogonal variable. **(F)** CDFs for decoding orthogonal values across vowels. Asterisks show significant differences between vowel and orthogonal (A-D, rank-sum or sign-rank depending on pairing, *p* < 0.05) or across orthogonal variables (Kruskal-Wallis, *p* < 0.05).

**Fig S5.**
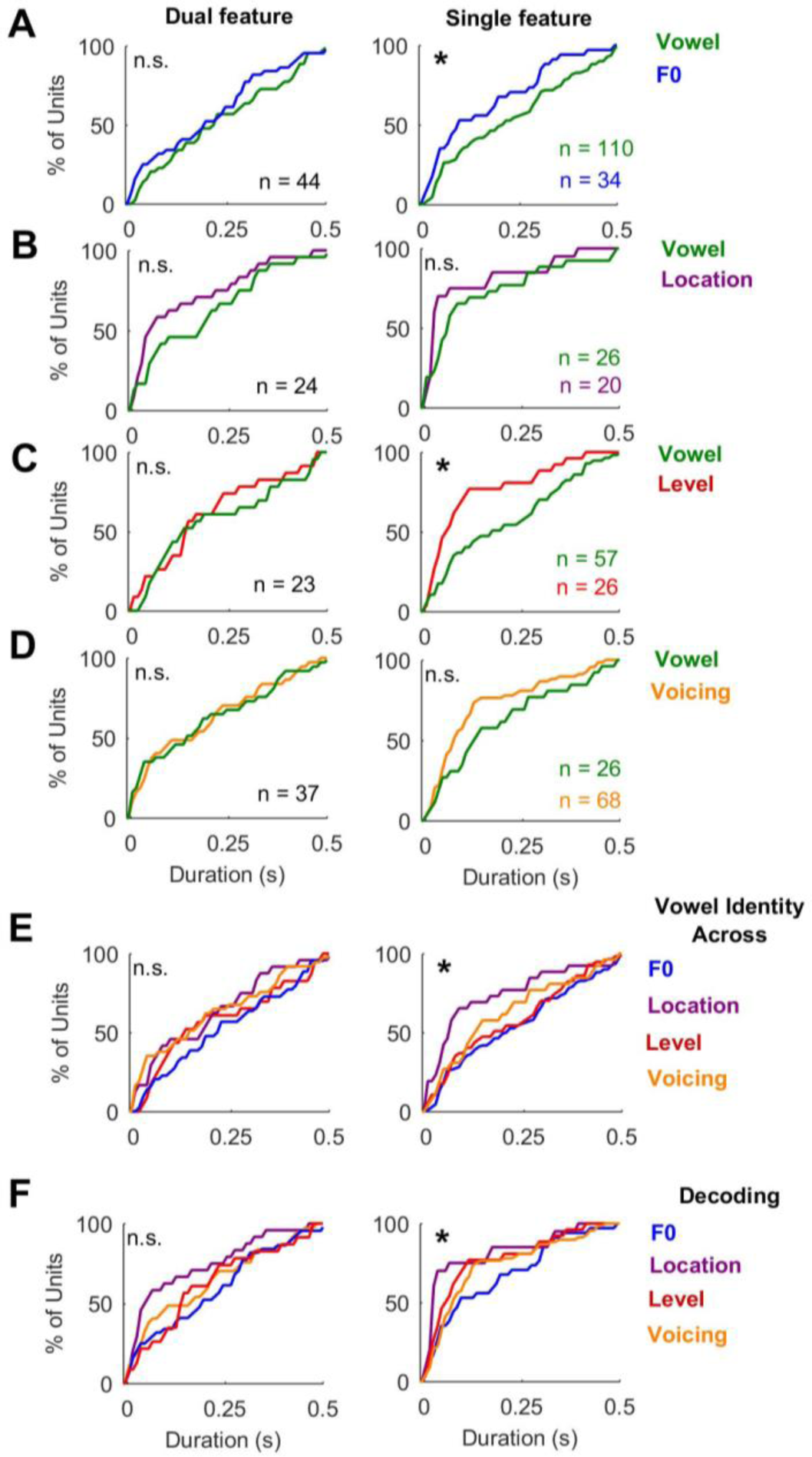
Temporal profiles for the duration of best decoding windows. **(A-D)** Cumulative distributions showing duration for best performance when decoding vowel identity or orthogonal variables (**A:** F0, **B**: location, **C**: level and **D**: voicing). Units are shown separately by classification as informative about vowel identity and orthogonal values (Dual feature units), or only vowel identity or orthogonal values (Single feature units). **(E)** CDFs for decoding vowel identity across each orthogonal variable. **(F)** CDFs for decoding orthogonal values across vowels. Asterisks show significant differences between vowel and orthogonal (A-D, rank-sum or sign-rank depending on pairing, *p* < 0.05) or across orthogonal variables (Kruskal-Wallis, *p* < 0.05).

**Fig S6.**
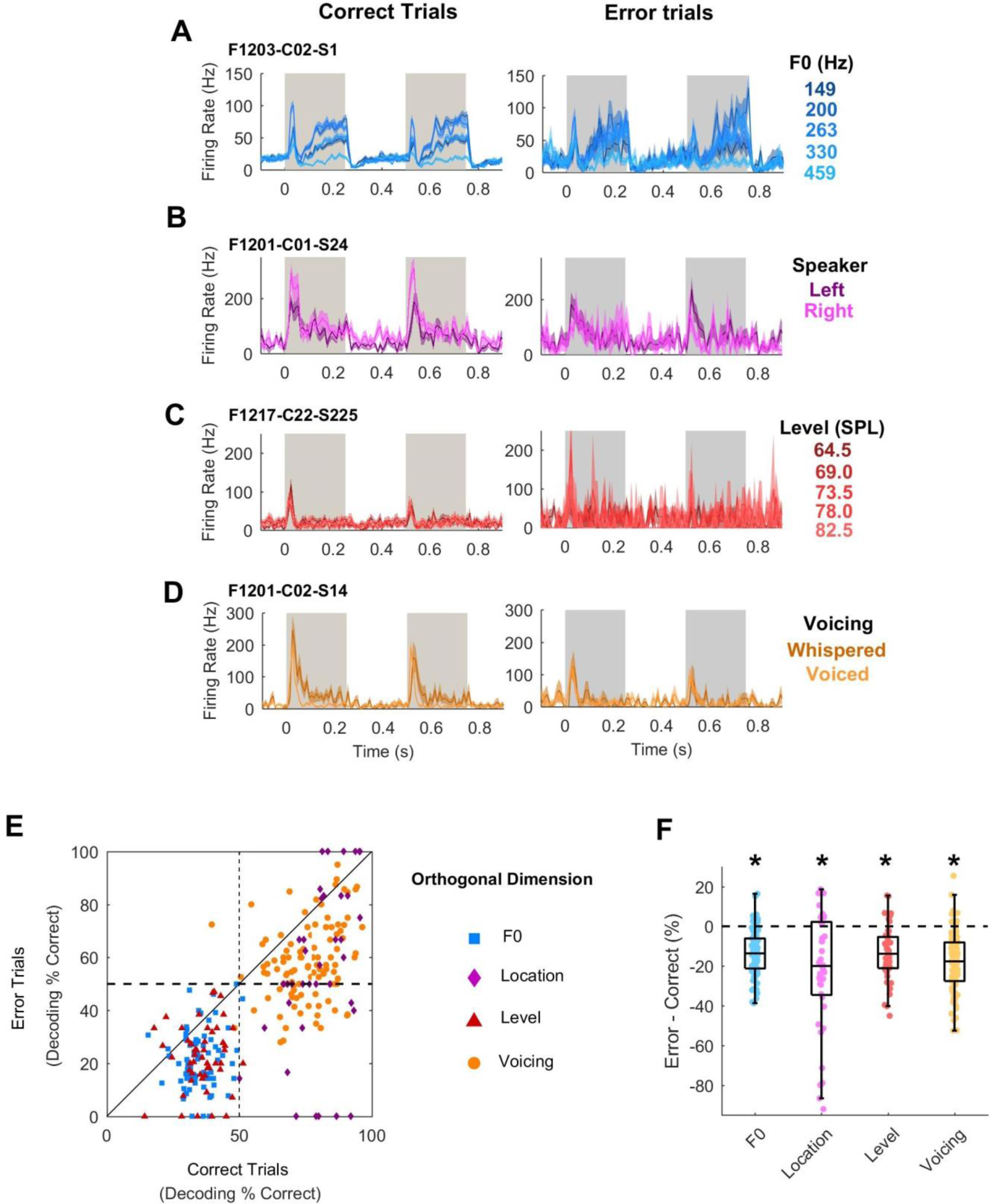
Encoding of orthogonal variables on correct and incorrect trials. (**A-D**) Example units show encoding of orthogonal variables on correct and error trials. Data is shown as mean ± s.e.m. (**E-F**) Change in decoding performance from correct to error trials. Asterisks show significant comparisons (paired t-test: F0: t_77_ = -10.3, *p* = 3.42 x 10^-16^, across location: t_40_ = -4.82, *p* = 2.12 x 10^-5^, across sound level: t_48_ = -6.98, *p* = 7.85 x 10^-9^, across voicing: t_104_ = -11.9, *p* = 4.52 x 10^-21^)

**Fig S7.**
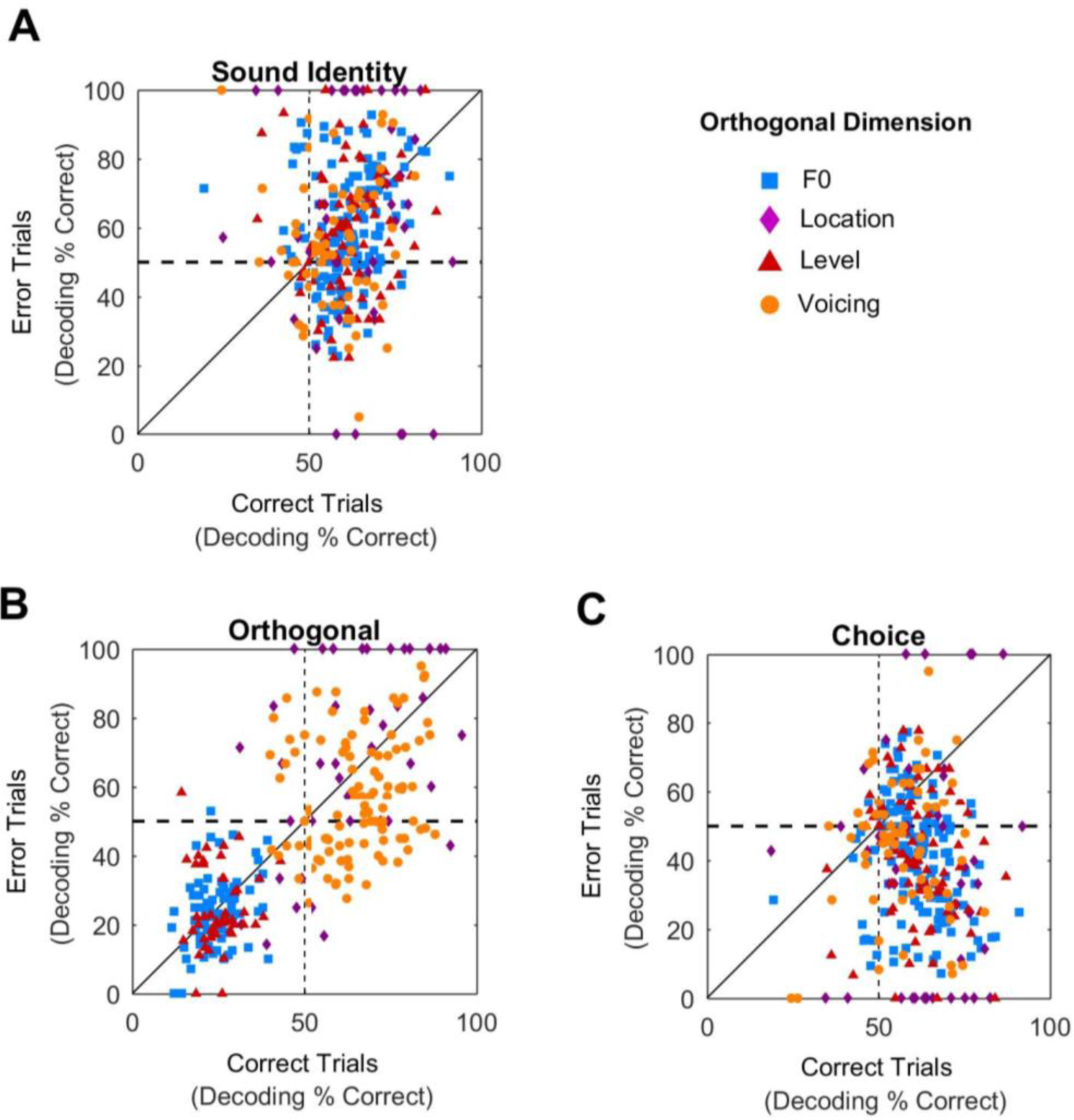
Error trial decoding performance using a fixed decoding window. Decoding of vowel identity (**A**), orthogonal values such as F0 or sound level (**B**) and behavioral response (**C**) on correct and error trials using a fixed time window in the first stimulus presentation (0 to 250 ms). There was no consistent difference between correct and error trials when decoding vowel identity or orthogonal values (*p* > 0.1 for all comparisons). However decoding the animal’s response direction was worse on error than correct trials (Across F0: t_153_ = -12.1, *p* = 3.40 x 10^-24^, across location: t_45_ = -5.5, *p* = 1.93 x 10^-6^, across sound level: t_79_ = -8.1, *p* = 5.77 x 10^-12^, across voicing: t_62_ = -4.7, *p* = 1.45 x 10^-5^). The effects of trial accuracy were greater on behavioral response than stimulus identity when sounds varied across F0 (t_306_ = 8.4, *p* = 2.15 x 10^-15^), across location (t_90_ = 3.8, *p* = 2.94 x 10^-4^), across sound level (t_158_ = 5.3, *p* = 3.90 x 10^-7^) and across voicing (t_124_ = 2.9, *p* = 0.004).

**Fig S8.**
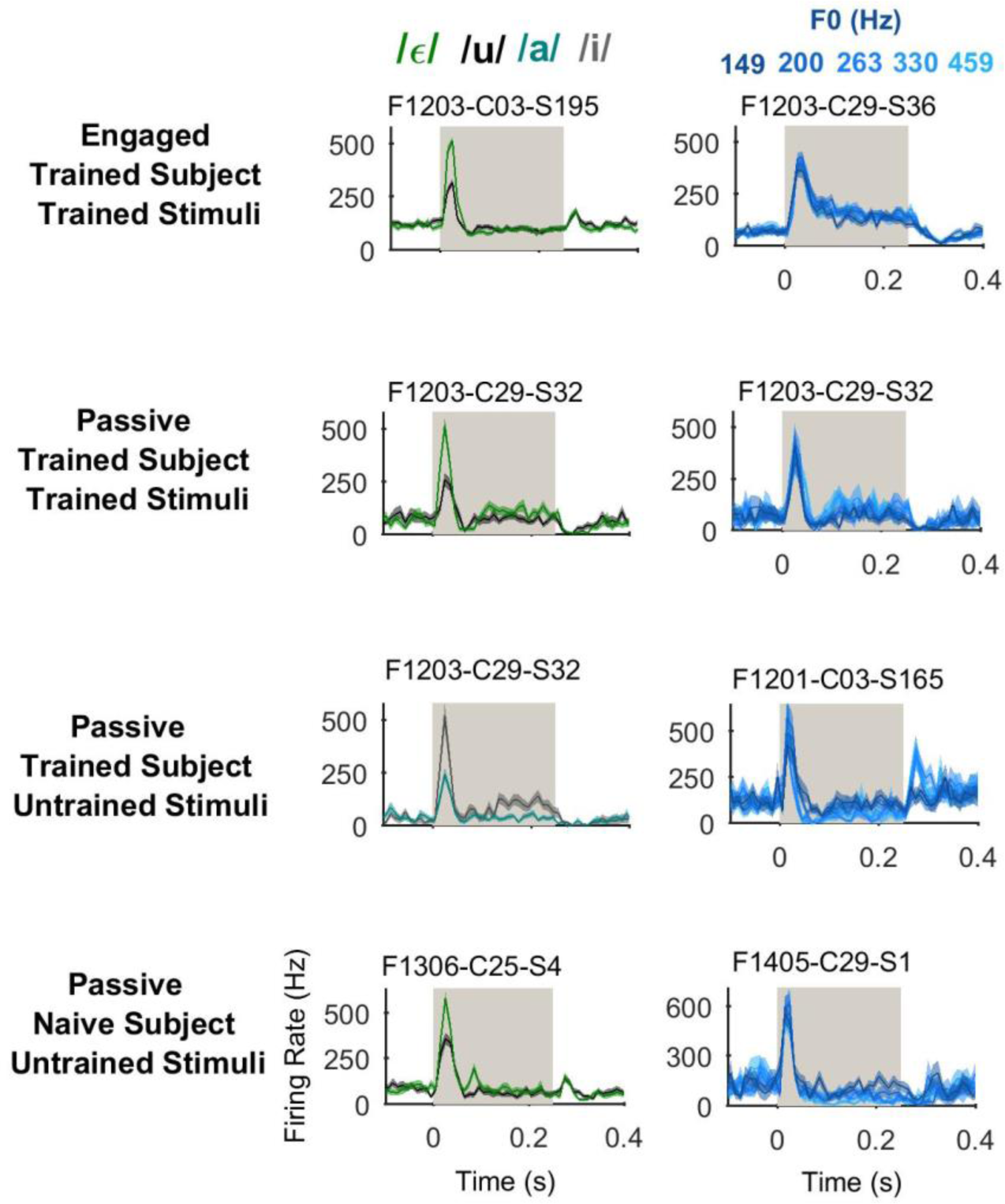
Example responses during task engagement and passive listening. Sound-evoked responses of individual unit examples, in task-engaged and passively listening, trained and untrained animals to trained and untrained vowels. Plots show mean ± s.e.m. firing rate.

**Fig S9.**
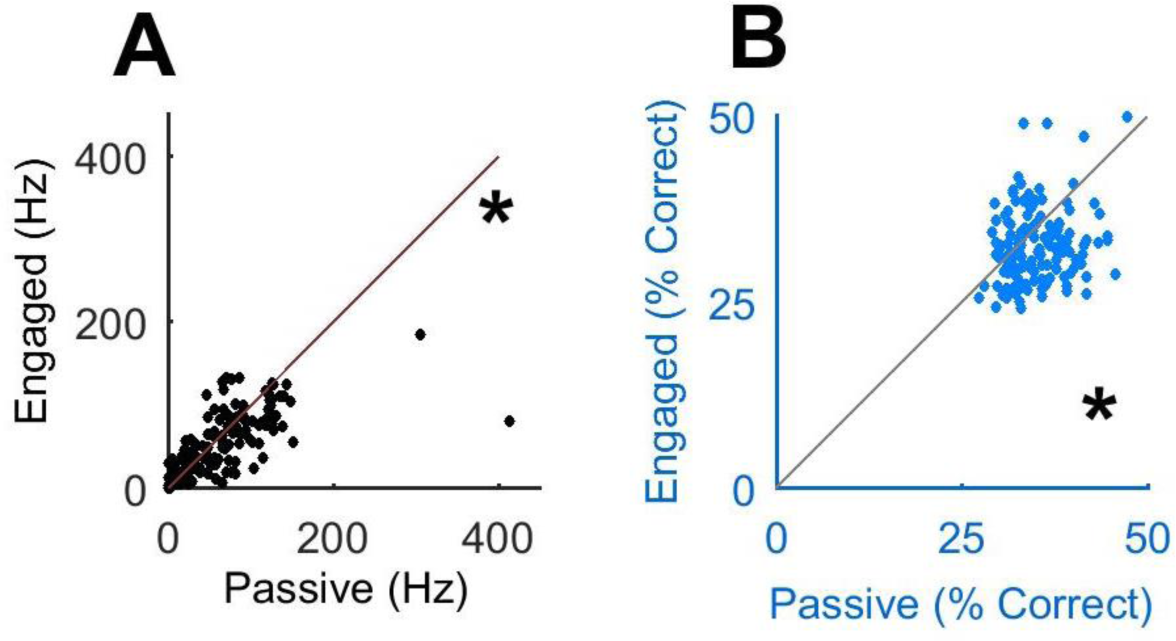
Example responses during task engagement and passive listening. (**A**) Firing rate in the time window that gave best performance decoding F0 (optimized independently for each unit in each experimental condition [passive/ engaged]). Data points indicate individual units (**B**) Paired comparison of best performance decoding F0 in optimized time windows. Data is shown as in (A). Asterisks show significant engagement-related suppression (*p* < 0.05).

**Fig S10.**
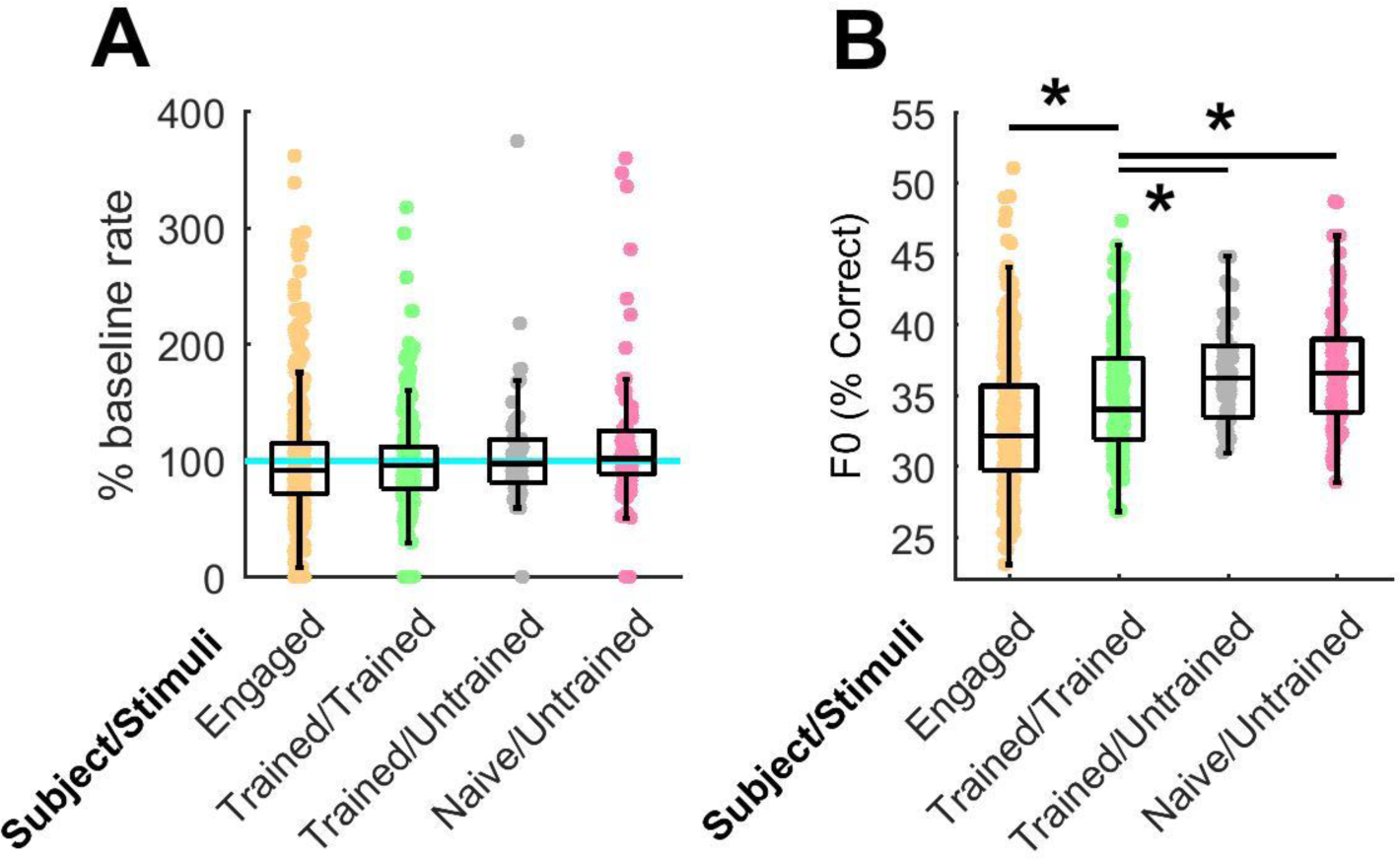
Effects of training on spiking activity and performance in time windows optimized for decoding F0. (**A-B**) Comparison of spiking activity normalized to baseline firing (A) and best performance decoding F0 (B) in optimized time window. Individual data points show individual units; box plots show median and inter-quartile ranges. Data also shown for task engaged responses for reference. Asterisks show significant comparisons between experimental groups: Normalized firing rates did not differ significantly between neurons recorded in any passive conditions, or between units responding to trained sounds during task engagement and passive listening. Decoding performance across all units differed significantly between groups (Kruskal-Wallis anova, χ^2^ = 21.0, *p* = 2.76 x 10^-5^) with decoding being significantly worse in units recorded from trained than naïve animals (Tukey-Kramer corrected for multiple comparisons, *p* = 1.0 x 10^-4^), and worse for units responding to trained than untrained sounds (*p* = 0.007). Performance decoding F0 of untrained sounds in naïve and trained animals was not significantly different (*p* = 0.935). Decoding performance of units responding to trained sounds during task engagement was significantly worse than when passively listening (*z* = 9.81, *p* = 1.06 x 10^-8^).

